# Revisiting the potency of Tbx2 expression in transforming outer hair cells into inner hair cells at multiple ages *in vivo*

**DOI:** 10.1101/2023.09.17.558159

**Authors:** Zhenghong Bi, Minhui Ren, Yu Zhang, Shunji He, Lei Song, Xiang Li, Zhiyong Liu

**Affiliations:** Institute of Neuroscience, State Key Laboratory of Neuroscience, CAS Center for Excellence in Brain Science and Intelligence Technology, Chinese Academy of Sciences, Shanghai, 200031, China; University of Chinese Academy of Sciences, Beijing, 100049, China; Department of Otolaryngology-Head and Neck Surgery, Ninth People’s Hospital, Shanghai Jiao Tong University School of Medicine, Shanghai, China; Ear Institute, Shanghai Jiao Tong University School of Medicine, Shanghai, China; Shanghai Center for Brain Science and Brain-Inspired Intelligence Technology, Shanghai, 201210, China

**Keywords:** Tbx2, Ikzf2, outer hair cell, inner hair cell, inner ear, cochlea

## Abstract

The cochlea, the auditory organ, contains two types of sound receptors: inner hair cells (IHCs) and outer hair cells (OHCs). Tbx2 is expressed in IHCs but repressed in OHCs. The neonatal OHCs with Tbx2 misexpression transdifferentiate into IHC-like cells. However, the extent of the switch from OHCs to IHC-like cells and the underlying molecular mechanism remain poorly understood. Furthermore, it is unknown whether Tbx2 can transform fully mature adult OHCs into IHC-like cells. In this study, we employ single-cell transcriptomic analysis, revealing that 85.6% of IHC genes, including *Slc17a8*, are upregulated, whereas only 38.6% of OHC genes, including *Ikzf2* and *Slc26a5*, are downregulated in neonatal OHCs with Tbx2 misexpression. Thus, our findings suggest that Tbx2 cannot fully reprogram neonatal OHCs into IHCs, contrary to previous assumptions. Consistently, Tbx2 also fails to fully reprogram cochlear progenitors into IHCs. Finally, Ikzf2 restoration alleviates the abnormalities present in the Tbx2+ OHCs, supporting the notion that Ikzf2 repression by Tbx2 contributes to the transdifferentiation of OHCs into IHC-like cells. Overall, our study reevaluates the effects of ectopic Tbx2 expression on OHC lineage development at different stages and provides molecular insights into how Tbx2 disrupts the gene expression profiles of OHCs. This research also lays the groundwork for future studies on OHC regeneration.

**Significance Statement:** Understanding the molecular and genetic mechanisms that govern the determination and stability of cochlear inner hair cells (IHCs) and outer hair cells (OHCs) would provide valuable insights into regenerating damaged IHCs and OHCs. In this manuscript, we conduct *in vivo* conditional overexpression of Tbx2 in cochlear sensory progenitors, neonatal OHCs and adult OHCs, respectively. Our findings challenge previous assumptions by revealing that Tbx2 expression alone can partially destabilize OHC fates but cannot fully convert OHCs into IHCs. Specifically, we demonstrate that the repression of Ikzf2 expression due to Tbx2 overexpression is one of the key pathways disrupting OHC fate.

## Introduction

Understanding how lineage specific or determining transcriptional factors (TFs), such as MyoD for muscle and Ascl1 for nerve, work is of importance in developmental biology field ^1^. It will provide new molecular and genetic insights of how each cell type is specified and maintained, with the possibility of opening avenues toward regenerating the missing cell population after damage. The mammalian auditory organ cochlea contains two subtypes of sound receptors that are known as hair cells (HCs), inner HCs (IHCs) and outer HCs (OHCs) ^2^. Prestin (*Slc26a5*) is expressed in OHCs but not in IHCs ^3–5^, and vGlut3 is opposite ^6,7^. Multiple subtypes of supporting cells (SCs) are intermingled with IHCs or OHCs ^8^. Both IHCs and OHCs are descendants of Atoh1+ cochlear prosensory progenitors ^9,10^. Atoh1 is a master TF essential for cochlear HC production ^11,12^. Nevertheless, Atoh1 should not be directly involved in the bifurcation of the progenitors into IHCs or OHCs, because both IHC and OHC development are severely defective in *Atoh1^-/-^* mice ^13–17^. Notably, although SCs are also derived from progenitors experiencing *Atoh1* mRNA ^9,10,18^, Atoh1 protein is never detectable in SCs, whereas strong Atoh1 protein is transiently expressed in IHCs and OHCs ^18^.

How the cochlear sensory progenitors differentiate into vGlut3+ IHCs or Prestin+ OHCs is recently unveiled by a series of genetic studies. Both Insm1 and Ikzf2 proteins are expressed in OHCs but not IHCs, and play key roles in relatively early and late OHC development, respectively ^19^. *Insm1^-/-^* OHCs transdifferentiate into IHC-like cells ^20,21^, and *Ikzf2^cello/cello^* OHCs are dysfunctional and maintain some IHC gene expression^22^. Moreover, ectopic Ikzf2 can induce Prestin expression in IHCs and promote regenerating Prestin+ OHC-like cells from adult SCs ^22,23^. Tbx2 is expressed as early as otocyst is formed ^24,25^. Recent three independent studies show that Tbx2 is essential in IHC fate specification, differentiation and maintenance ^26–28^. Briefly, *Tbx2^-/-^* IHCs transdifferentiate into OHC-like cells, however, there is a debate regarding the extent of cell fate conversion from IHCs to OHCs. While one study indicates that *Tbx2^-/-^* IHCs fully become OHCs ^27^, the other study supports that such a cell fate conversion is largely incomplete ^26^. Conversely, the neonatal OHCs misexpressing ectopic Tbx2 decrease Prestin and increase vGlut3 expression ^27,28^. Nonetheless, the degree to which the forced Tbx2 expression transforms OHCs into IHC-like cells remains poorly understood. Moreover, it is unknown whether the adult OHCs are responsive to Tbx2 misexpression. Neither is known about the molecular mechanisms underlying how Tbx2 converts OHCs to IHC-like cells.

In our current study, we addressed the three questions above. When Tbx2 was overexpressed in neonatal OHCs, the OHC to IHC conversion was incomplete. Approximately, 85.6% of IHC genes including vGlut3 were upregulated, in contrast only 38.6% of OHC genes such as Prestin and Ikzf2 were downregulated. To our surprise, we found that Tbx2 seemed unable to fully reprogram the cochlear progenitors into IHCs, either. Nonetheless, Tbx2 was competent to perturb the gene expressions of adult OHCs. Finally, we demonstrated that Ikzf2 restoration partially alleviated the abnormalities of the Tbx2+ OHCs. Thus, repressing Ikzf2 by Tbx2 should be one of the key cascades involved in switching OHCs to IHC-like cells.

## Results

### Generation of a new conditional *Tbx2* overexpression mouse line

To induce ectopic Tbx2 effectively and conditionally in different cell types or at distinct ages, we established a new *Rosa26*-Loxp-stop-Loxp-Tbx2*3ξHA-P2A-tdTomato (*Rosa26^Tbx^*^2^*^/+^* in brief). In the *Rosa26^Tbx^*^2^*^/+^*, Tbx2 and tdTomato expressions were tightly paired by the 2A strategy (Supplemental Figure 1A-C). Southern blot confirmed that there was no random insertion of the targeting vector in mouse genome (Supplemental Figure 1D). The wild type (WT), heterozygous knockin (KI) *Rosa26^Tbx^*^2^*^/+^* and homozygous (*Rosa26^Tbx^*^2^*^/Tbx^*^2^) were healthy, fertile, and were readily to be identified by tail PCR (Supplemental Figure 1E). *Rosa26^Tbx^*^2^*^/+^* and *Rosa26^Tbx2/Tbx2^*did not exhibit obvious developmental defect.

According to our design above, upon the Cre-mediated recombination, in principle cells expressing Tbx2, which was tagged with three HA fragments at its C-terminus, were also permanently traced by tdTomato. The tdTomato expression would certainly facilitate subsequent fate mapping and transcriptomic analysis at the single-cell resolution. As described below, we would use *Slc26a5^CreER/+^* to specifically target cochlear OHCs at both postnatal day 2 (P2)/P3 and P60/P61, respectively, as well as *Atoh1^Cre/+^* to target the undifferentiated cochlear sensory progenitors at embryonic ages.

### Ectopic Tbx2 converts neonatal OHCs into IHC-like cells

To demonstrate whether functional Tbx2 could be effectively induced in *Rosa26^Tbx2/+^,* we overexpressed Tbx2 in neonatal cochlear OHCs by crossing it with *Slc26a5^CreER/+^* ^29^. It is known that neonatal OHCs misexpressing Tbx2 decrease Prestin and increase vGlut3 ^27,28^. Given the effective Tbx2 induction, we should observe the similar phenotype in *Slc26a5^CreER/+^*; *Rosa26^Tbx^*^2^*^/+^* mice (Slc26a5-Tbx2 in brief), but not in the control *Slc26a5^CreER/+^*; *Rosa26*-CAG-Loxp-stop-Loxp-tdTomato (Ai9)/+ (Slc26a5-Ai9 in brief). In Slc26a5-Ai9 cochleae, OHCs were equivalent to wild type OHCs, except expressing tdTomato. Both Slc26a5-Ai9 and Slc26a5-Tbx2 mice were administered tamoxifen at P2/P3, and analyzed at P14 or P42 or P76 or P120 (Figure 1A, n=3 at each age). In control Slc26a5-Ai9, almost all OHCs were tdTomato+ regardless of when they were analyzed. Thus, the data at P42 were chose to present as examples. Three rows of Prestin+ OHCs and one row of vGlut3+ IHCs were regularly aligned at P42 (Figure 1B-B’’’). Almost all OHCs were tdTomato+, but IHCs never expressed tdTomato, in agreement with the OHC exclusive Cre activity of *Slc26a5^CreER/+^* ^29^.

**Figure 1.**
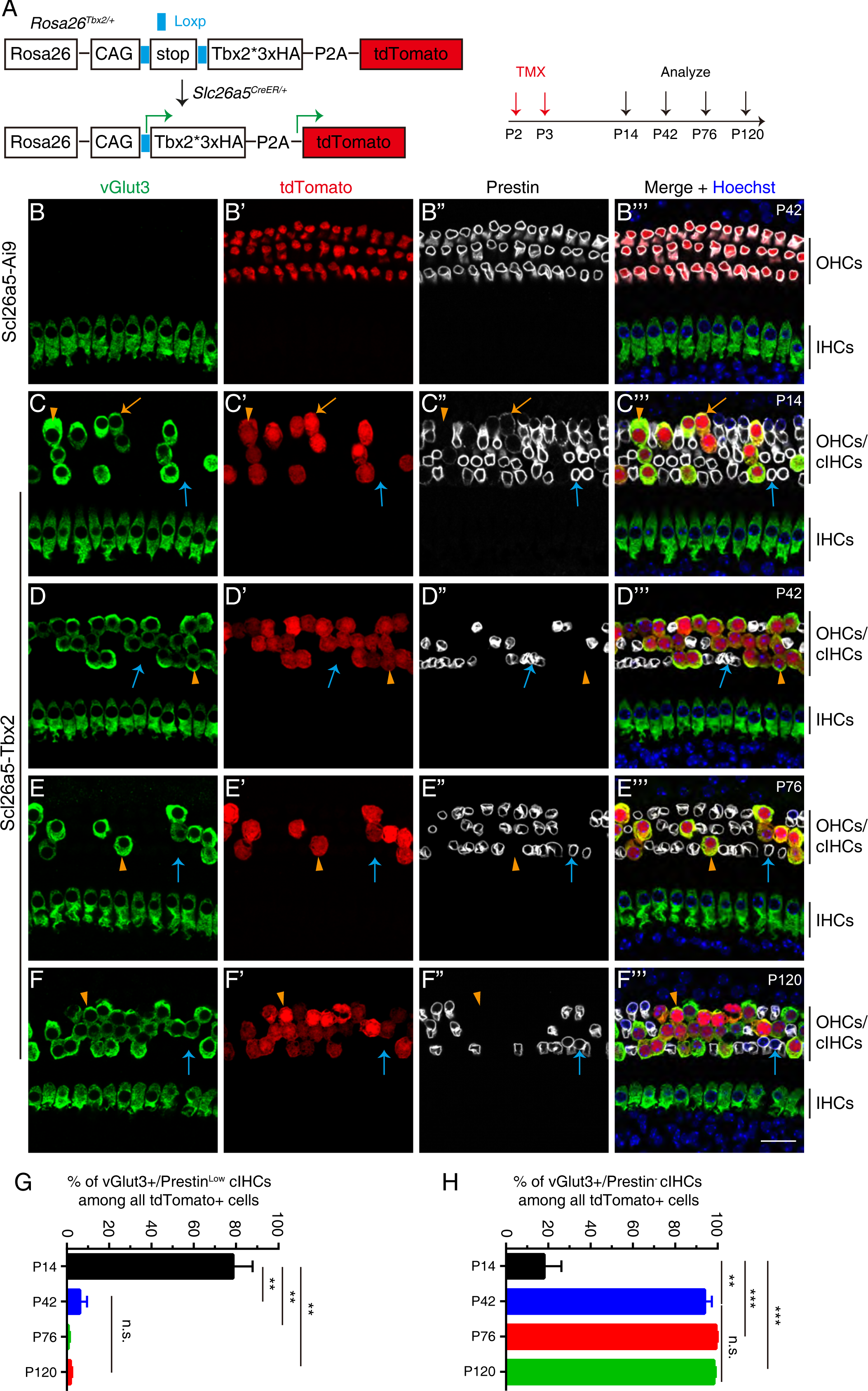
vGlut3 is reactivated and Prestin is repressed when Tbx2 is ectopically induced in neonatal OHCs. **(A)** A simple illustration of how Tbx2 is specifically overexpressed in neonatal OHCs which normally do not express Tbx2. **(B)** Triple staining of vGlut3, tdTomato and Prestin in mice with different genotypes: Slc26a5-Ai9 at P42 (B-B’’’) and Slc26a5-Tbx2 at four different ages, P14 (C-C’’’), P42 (D-D’’’), P76 (E-E’’’) and P120 (F-F’’’). The images were taken from the apical turns where the cell death was minimal. The orange arrows mark the cIHCs that express vGlut3 and low level of Prestin, whereas orange arrowheads label the vGlut3+ cIHCs in which Prestin is undetectable. The blue arrows point to the endogenous OHCs that maintain high level of Prestin and do not express tdTomato (Tbx2) and vGlut3. **(G-H)** Quantification of the percentage of the cIHCs expressing high vGlut3 and low Prestin (G), and the cIHCs expressing high vGlut3 but no detectable Prestin (H). Data are presented as Mean ± Sem (n=3 for each age). Student’s *t* test was used for statistical analysis. ** p<0.01; ***p<0.001; n.s., not significant. OHCs: outer hair cells; IHCs: inner hair cells; cIHCs: converted inner hair cells. Scale bar: 20 μm (F’’’).

In contrast, high level of vGlut3 was expressed, but Prestin was marked decreased (orange arrows in Figure 1C-C’’’) or became undetectable (orange arrowheads in Figure 1C-C’’’) in all tdTomato (Tbx2)+ OHCs in the Slc26a5-Tbx2 at P14. According to our criteria, those vGlut3+/tdTomato+ OHCs expressing low (Prestin^Low^) or no detectable Prestin (Prestin^-^) were defined as converted IHCs (cIHCs in brief) derived from the endogenous OHCs. The heterogenous levels of Prestin in the cIHCs indicated the various stages of incomplete conversion from OHCs to IHCs or IHC-like cells, with Prestin^-^ cIHCs representing the highest degree of conversion. Notably, the endogenous OHCs not expressing tdTomato (Tbx2) maintained the Prestin expression at a level as high as the OHCs in control mice and as expected did not express vGlut3 (blue arrows in Figure 1C-C’’’). Moreover, all vGlut3+ cIHCs were tdTomato (Tbx2)+ and vice versa in Slc26a5-Tbx2. Notably, despite the identical tamoxifen treatment, numbers of the tdTomato+ cells in Slc26a5-Tbx2 were less than in Slc26a5-Ai9 mice. It was likely due to that the long polycistronic Tbx2/Tdtomato transcription in *Rosa26^Tbx^*^2^*^/+^* was not as efficient as the Tdtomato alone in Ai9/+ mice. Collectively, our data show that the Slc26a5-Tbx2 mouse line is an efficient model to induce cIHCs from neonatal OHCs.

### HC loss occurs in the Slc26a5-Tbx2 mice after P14

The long-term effects of Tbx2 misexpression on OHCs remains unknown. We further analyzed the Slc26a5-Tbx2 mice at P42 (Figure 1D-D’’’), at P76 (Figure 1E-E’’’) or at P120 (Figure 1F-F’’’). Briefly, similar phenotypes of cell fate conversion from OHCs to IHCs were observed. In control mice at P120 (n=3), HCs were largely intact with sporadic death primarily in the basal turn (Supplemental Figure 2A-C). In contrast, after P14 we started to observe loss of cIHCs or the endogenous OHCs in a base to apex gradient in Slc26a5-Tbx2 mice. Notably, the death of the endogenous OHCs not expressing Tbx2 should be a secondary effect due to degeneration of nearby cIHCs. At P42, the cIHCs/OHC loss primarily occurred in basal turn (Supplemental Figure 2D-F). Approximately, 282.67 ± 76.17 (n=3) cells (Prestin+ endogenous OHCs and vGlut3+ cIHCs) existed in basal turn of Slc26a5-Tbx2 mice at P42, significantly (*p<0.05) less than the 732.33 ± 107.20 (n=3) Prestin+ OHCs in basal turn of Slc26a5-Ai9 mice (Supplemental Figure 2J).

We noticed that the cell death not only migrated into middle turn, but also became more severe in basal and middle turn in Slc26a5-Tbx2 cochleae at P120 (Supplemental Figure 2G and H). Notably, the cell death in apical turn was very mild in Slc26a5-Tbx2 mice at P120 (Supplemental Figure 2I). By grouping basal and middle turns together, we found that 226.30 ± 34.37 (Prestin+ endogenous OHCs and vGlut3+ cIHCs) were present in Slc26a5-Tbx2 mice at P120 (n=3), which were much less (*** p<0.001) than the 1300 ± 71.16 (n=3) Prestin+ OHCs in Slc26a5-Ai9 mice (Supplemental Figure 2K). The IHCs in all three turns were normal at P42. In contrast, severe IHC death happened in basal and middle turns, but not in apical turn, at P120, which again was likely due to the secondary effect of cIHCs/OHC loss. In sum, those cIHCs permanently expressing Tbx2 were unhealthy and started to degenerate after P14.

### The cell fate switch from OHCs to cIHCs is largely complete by P42

Next, we determined when the cell fate switch from OHCs to cIHCs was complete after Tbx2 was induced in OHCs by quantifying the percentages of vGlut3+/Prestin^Low^ cIHCs (orange arrows in Figure 1) or vGlut3+/Prestin^-^ cIHCs (orange arrowheads in Figure 1), respectively, at P14, P42, P76 and P120. Due to the none or limited cIHCs/OHCs loss in apical turns after P14, to simply the analysis, we focused on the apical turns of the Slc26a5-Tbx2 mice. Thus, expression of vGlut3 and absence of Prestin in the cIHCs suggest that conversion of OHCs into cIHCs is completed by P42. Among all tdTomato+ cIHCs, the percentage of vGlut3+/Prestin^Low^ cIHCs was 78.52% ± 9.27% at P14 (n=3), markedly (**p<0.01) dropped to 6.12% ± 3.40% at P42 (n=3), and nearly disappeared at P76 (n=3) and at P120 (n=3) (Figure 1G). Conversely, the percentage of vGlut3+/Prestin^-^ cIHCs was 18.04% ± 8.17% at P14, dramatically (**p<0.01) increased to 93.88% ± 3.40% at P42, and became nearly ∼100% at P76 and at P120 (Figure 1H). Collectively, we concluded that, upon Tbx2 induction in neonatal OHCs, vGlut3 upregualtion was rapid and all cIHCs expressed high level of vGlut3 by P14, whereas Prestin downregulation was gradual and a large fraction of cIHCs belonged to Prestin^Low^ by P14. Thus, the cell fate conversion process of the cIHCs was not synchronized at P14, but largely completed by P42.

### The cIHCs also gain other IHC features and lose Ikzf2 expression

We next determined whether cIHCs expressed other IHC markers or gain additional IHC properties at P42. Triple staining of adult IHC marker Slc7a14, tdTomato and Ctbp2 showed that, different from the control mice (Supplemental Figure 3A-A’’’), besides the IHCs (#1, blue arrowheads in Supplemental Figure 3B-B’’’), Slc7a14 was also weakly expressed in the tdTomato+ cIHCs (#2, orange arrows in Supplemental Figure 3B-B’’’). Notably, the endogenous OHCs not expressing tdTomato did not express Slc7a14 (#3, blue arrows in Supplemental Figure 3B-B’’’). Moreover, the endogenous OHCs (#3) did not express Ctbp2 in their nuclei, however, the nuclear Ctbp2 expression level in the cIHCs (#2) was as high as that in the endogenous IHCs (#1).

We further measured the nuclei diameter and quantified the number of Ctbp2+ puncta of the IHCs (#1, 314 cells), cIHCs (#2, 296 cells) and the endogenous OHCs (#3, 385 cells) in the same region of the Slc26a5-Tbx2 mice (n=3). The nuclei size of the cIHCs reached the size of the IHCs and was significantly (***p<0.001) larger than the OHCs (Supplemental Figure 3C). However, the number of Ctbp2+ puncta (2.54 ± 0.34) in the cIHCs was still significantly (****p<0.0001) less than 16.24 ± 0.12 in the IHCs (Supplemental Figure 3D). We also checked the expression pattern of another IHC marker Otoferlin that was only expressed in IHCs in control mice at P42 (Supplemental Figure 3E-E’’). As expected, Otoferlin was expressed in the tdTomato+ cIHCs (orange arrows in Supplemental Figure 3F-F’’), but was absent in the endogenous OHCs not expressing tdTomato (blue arrows in Supplemental Figure 3F-F’’).

We wondered whether Ikzf2, a key TF for stabilizing OHC fate ^22^, was repressed in the cIHCs by taking advantage of the *Ikzf2^V^*^5^*^/+^*mice in which Ikzf2 is tagged with three V5 tags at its C-terminus ^21,26^. V5 (Ikzf2) was specifically expressed in all OHCs in control *Ikzf2^V^*^5^*^/+^*mice (Supplemental Figure 3G-G’’). However, V5 (Ikzf2) expression disappeared in the vGlut3+ cIHCs in the *Slc26a5^CreER/+^*; *Rosa26^Tbx2/+^*; *Ikzf2^V5/+^* (Slc26a5-Tbx2-*Ikzf2^V5/+^* in brief) mice (orange arrows in Supplemental Figure 3H-H’’). In contrast, the endogenous OHCs not expressing vGlut3 maintained V5 (Ikzf2) expression (blue arrows in Supplemental Figure 3H-H’’). Collectively, the data above further supported that the cIHCs gained, albeit not completely, IHC features, and lost OHC characteristics, especially the Ikzf2 expression.

### The cIHCs lose electromotility and the hearing capacity is severely impaired in Slc26a5-Tbx2 mice at P42

We applied patch-clamp to three cell types at P42: 1) WT_OHCs (#1, n=6, cell number) from wild type mice; 2) endogenous OHCs (#2, n=11) not expressing tdTomato and 3) the tdTomato+ cIHCs (#3, n=9) in the same Slc26a5-Tbx2 mice (Figure 2A). Relative to two kinds of OHCs (#1 green and #2 blue), the non-linear capacitance (NLC) disappeared in the cIHCs (#3, red) (Figure 2A). The calculated total Prestin amount in the cIHCs were also significantly (**** p<0.0001) reduced, relative to the OHCs (Figure 2B). It agreed with that Prestin expression was markedly decreased or completely lost by immunostaining assay (Figure 1). We also noticed that the NLC or Prestin level in the OHCs (#2, blue) was slightly (*p<0.05) smaller than that in WT_OHCs (#1, green). It might be due to that the endogenous OHCs (#2) were, to some extent, affected by the nearby cIHCs (#3), although they maintained the OHC fates and maintain high Prestin expression (Figure 1). Moreover, we calculated the total cell surface area of the cIHCs by measuring the linear capacitance (Figure 2C). No statistic difference existed about cell surface area between WT_OHCs (#1) and OHCs (#2) (Figure 2C). It is known that IHCs are bigger and have larger cell surface area than OHCs ^30^. Therefore, we predicted that the cell surface of cIHCs would be larger than the OHCs. To our surprise, the total cell surface area of cIHCs (#3) was significantly (*p<0.05) smaller than that of OHCs (#2) (Figure 2C), albeit the nuclei size of the cIHCs was larger than that of OHCs (Supplemental Figure 3C).

**Figure 2.**
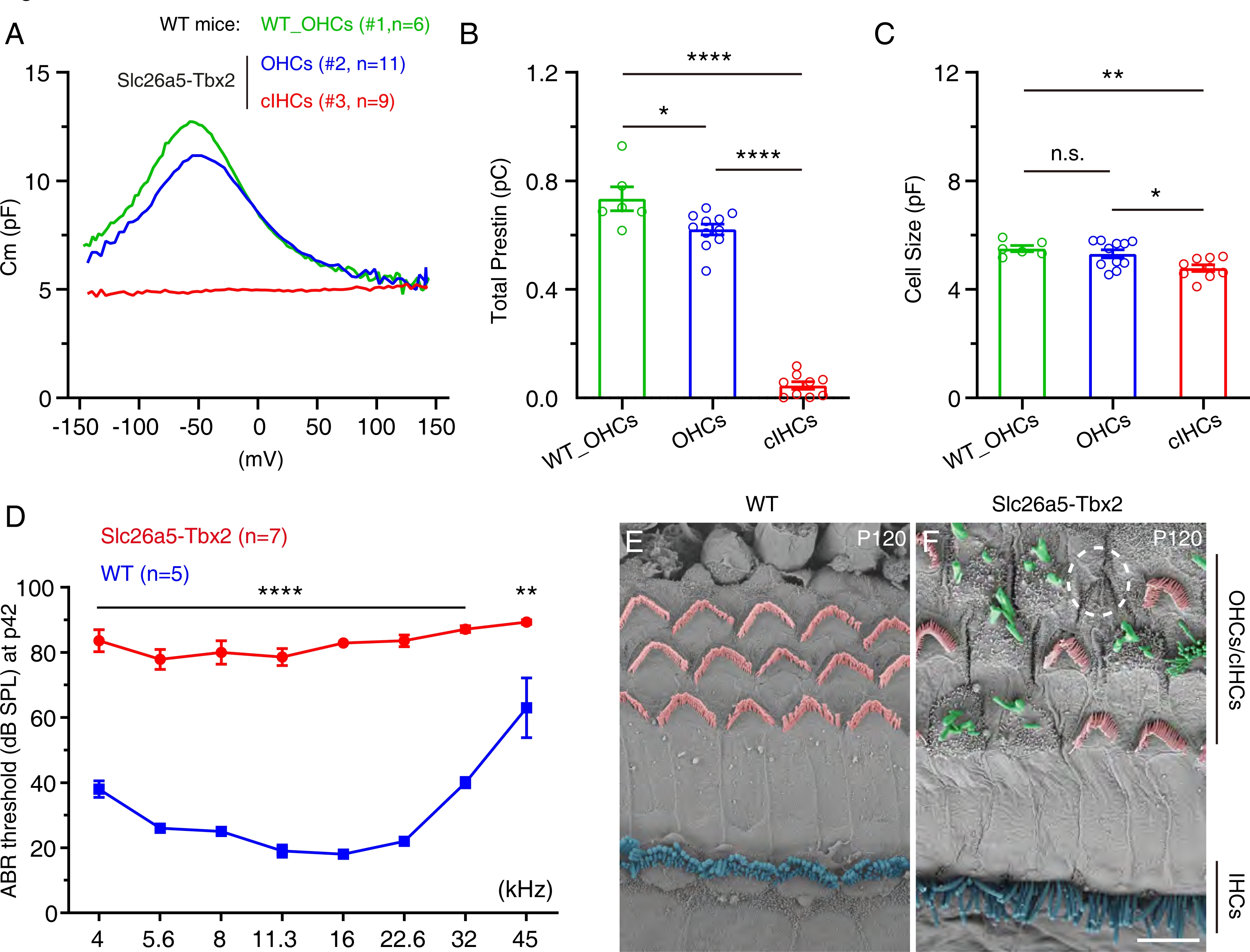
The cIHCs lose electromotility and their hair bundles are degenerated. **(A)** Non-linear capacitance (NLC) measurement of the OHCs in wild type (WT) mice at P42 (#1, green, n=6, cell number), and the endogenous OHCs not expressing tdTomato/Tbx2 (#2, blue, n=11) and the cIHCs expressing tdTomato/Tbx2 (#3, red, n=9). Notably, #2 and #3 are from the same Slc26a5-Tbx2 mice at P42, and they are readily to be distinguished by presence or absence of tdTomato. **(B-C)** Comparing the total Prestin amount (B) and cell size (C) among the three cell types in (A). Data are presented as Mean ± Sem. Statistical analysis was performed using one-way ANOVA. *p<0.05, ** p<0.01; ****p<0.0001; n.s., not significant. **(D)** ABR measurement of WT (n=5) and Slc26a5-Tbx2 (n=7) mice at P42. Significant hearing impairment occurs in Slc26a5-Tbx2 mice at all tested frequencies. Data are presented as Mean ± Sem. Student’s *t* test was used for statistical analysis. ** p<0.01; ****p<0.0001. **(E-F)** SEM analysis of WT (n=3) and Slc26a5-Tbx2 (n=3) mice at P120. The degenerated hair bundles of the cIHCs are shown in green color, whereas the white dotted circle marks one cIHC that either has died or lost hair bundles. Scale bars: 5 μm (F).

Consistently, auditory brainstem response (ABR) measurement showed that the hearing thresholds in all tested frequencies of Slc26a5-Tbx2 (n=7) were significantly (****p<0.0001 or **p<0.01) increased, relative to the wild type (n=5) mice (Figure 2D), meaning that the hearing capacity was severely impaired at P42. Finally, scanning electron microscope (SEM) assay demonstrated that, relative to the hair bundles of WT_OHCs in wild type mice (pink color in Figure 2E), the hair bundles of the cIHCs were degenerated (green color in Figure 2F) or completely lost (white dotted circle in Figure 2F) in Slc26a5-Tbx2 at P120. Notably, the hair bundles of the OHCs in Slc26a5-Tbx2 (pink color in Figure 2F) seemed largely normal. Collectively, Tbx2 misexpression converted neonatal OHCs to cIHCs that lost Prestin-mediated electromotility and displayed degenerated hair bundle, resulting in severe hearing impairment.

### The significantly different genes between wild type OHCs and IHCs at adult ages

Our analysis above supported that the cIHCs were not equivalent to IHCs. To further solidify this notion, we determined the degree of cell fate conversion in cIHCs by single cell transcriptomic profiling of the cIHCs. We have reported single cell RNA-seq profile of 17 wild type OHCs at P30 (P30_WT OHCs-old) ^23^ and the 50 wild type IHCs at P30 (P30_WT_IHCs) ^26^. To balance the cell numbers of IHCs and OHCs, we additionally picked 32 wild type OHCs at P30 (P30_WT OHCs-new) from Slc26a5-Ai9 mice. By mixing all IHCs and OHCs, as expected we found that new and old P30_WT OHCs were overlapped (Supplemental Figure 4A), thus the new and old P30_WT OHCs were grouped together as P30_WT OHCs. Notably, gene profiling of all cells was conducted by smart-seq approach which can yield high gene coverage because it is full-length RNA-seq ^31^.

Both P30_WT OHCs and P30_WT IHCs shared pan-HC markers *Pou4f3* and *Myo7a* (Supplemental Figure 4B and C) ^32^. Moreover, IHC markers *Slc17a8*, *Slc7a14* and *Tbx2* were enriched in P30_WT_IHCs (Supplemental Figure 4D-F) ^6,7,26,33^, whereas OHC markers *Slc26a5*, *Ikzf2*, and *Lbh* were enriched in P30_WT_OHCs (Supplemental Figure 4G-I) ^3,4,22,34^. We thoroughly compared the transcriptomic profiling between P30_WT OHCs and P30_WT IHCs and found that 2010 genes showed significantly (absolute value of fold change (FC)>4, and p<0.05) different levels, with 1482 genes enriched in P30_WT IHCs and 528 genes enriched in P30_WT OHCs (Supplemental table 1), which were defined as IHC or OHC genes, respectively. The top differently expressed genes were presented in heatmap (Supplemental Figure 4J).

### The cell fate conversion in cIHCs is largely incomplete

With the same smart-seq approach, we also manually picked 67 tdTomato+ cIHCs from the Slc26a5-Tbx2 mice at P42, which were defined as P42_cIHCs. We picked them at P42, instead of P30, in order to give them longer time to transform because they were derived from the endogenous OHCs at P2/P3, rather than the control OHCs that start differentiation from embryonic ages. P42_cIHCs were mixed with P30_WT IHCs, and P30_WT OHCs. Although the P42_cIHCs were closer to P30_WT IHCs than to P30_WT OHCs, three distinct cell clusters formed (Figure 3A and B). Nevertheless, the P42_cIHCs expressed pan-HC markers (*Myo7a*, *Pou4f3* and *Tmc1*), IHC markers (*Slc17a8*, *Slc7a14* and *Tbx2*), but not or minimally expressed OHC markers (*Slc26a5*, *Ikzf2* and *Lbh*) (Supplemental Figure 5A-C). It was worthy of pointing out that *Tbx2* levels seemed higher in P42_cIHCs than in P30_WT IHCs (Supplemental Figure 5C). It might be due to that both ectopic (from *Rosa26* locus) and the endogenous *Tbx2* were transcribed in P42_cIHCs. Moreover, the global gene expression correlation coefficient analysis showed that P42_cIHCs resembled P30_WT IHCs more than P30_WT OHCs, but manifest difference existed between P42_cIHCs and P30_WT IHCs (Supplemental Figure 5D).

**Figure 3.**
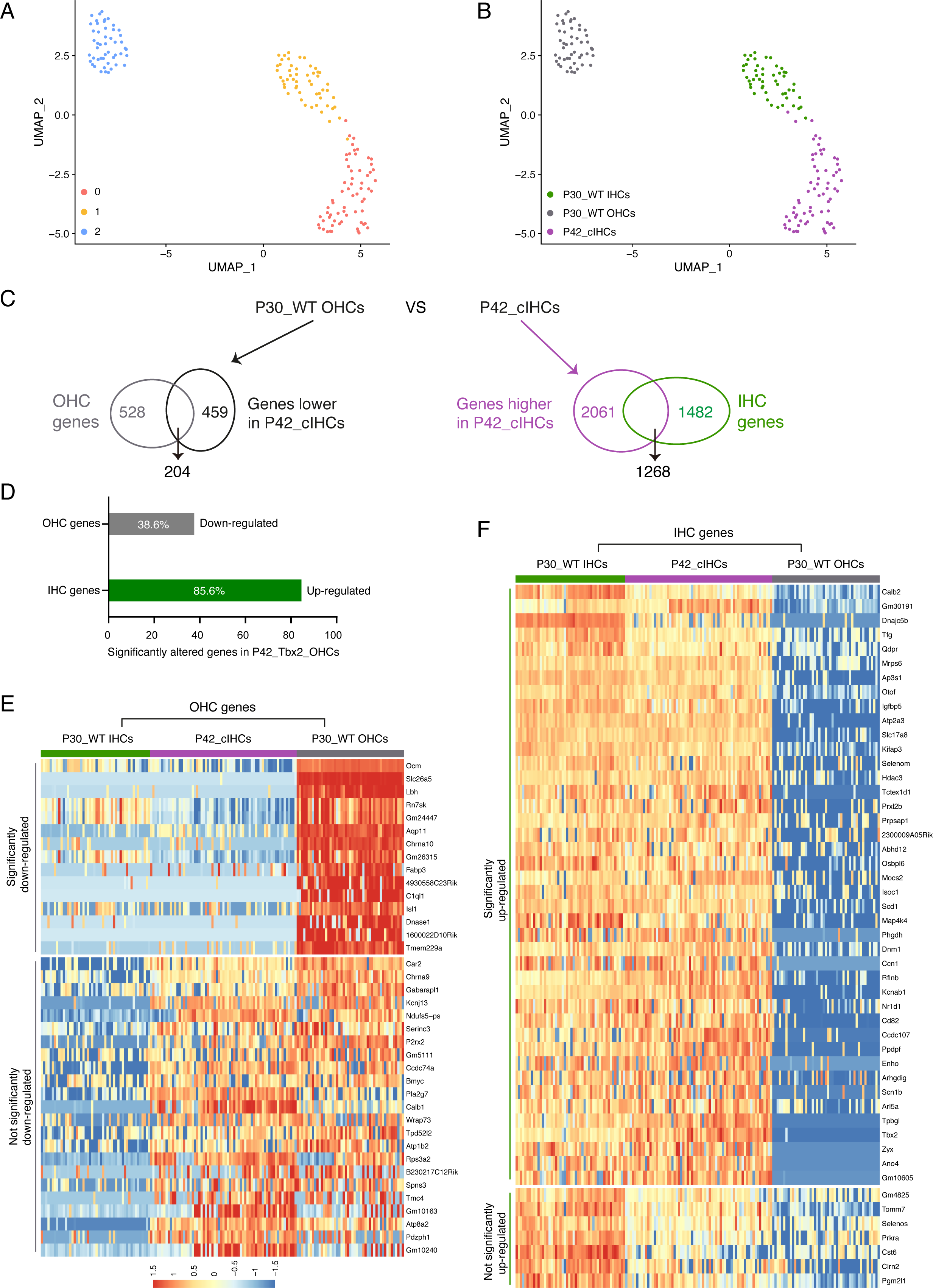
Single-cell transcriptomic analysis of wild type IHCs, OHCs and cIHCs at adult ages. **(A-B)** The cell population, which are mixed by P30_WT IHCs, P30_WT OHCs and P42_cIHCs, are subjected to unsupervised cluster (A) or marked by their own identities (B). **(C-D)** Transcriptomic comparison between P42_cIHCs and P30_WT OHCs reveals 459 genes that are expressed at a significantly lower level in P42_cIHCs than in P30_WT OHCs, among which 204 are the defined OHC genes (left side in C). Conversely, 2061 genes show opposite patterns and are expressed in a higher level in P42_cIHCs than in P30_WT OHCs, among which 1268 are the defined IHC genes (right side in C). Thus, relative to P30_WT OHCs, 38.6% of OHC genes, and 85.6% of IHC genes are significantly decreased or increased in P42_cIHCs, respectively (D). **(E)** Heatmap of the top significantly (upper part) or not significantly (lower part) downregulated OHC genes. **(F)** Heatmap of the top significantly (upper part) or not significantly (lower part) upregulated IHC genes.

To estimate how far the cIHCs were divergent from the OHCs, we compared the global gene profiles between P42_cIHCs and P30_WT OHCs. The expression levels of 2061 genes were dramatically (FC>4, and p<0.05) higher in P42_cIHCs than in P30_WT OHCs, whereas 459 genes exhibited the opposite (Figure 3C). There were 1268 genes overlapped between the 2061 genes and the 1482 IHC specific genes defined above (Supplemental table 1). Likewise, among the 459 genes, 204 were overlapped with the 528 OHC genes (Supplemental table 1). Thus, 85.6% (1268/1482) of IHC genes were markedly increased, and 38.6% (204/528) of OHC genes were significantly decreased in the P42_cIHCs (Figure 3C). The top significantly or not significantly altered OHC and IHC genes were presented in heatmap (Figure 3E and F) and all the gene list was included in Supplemental table 2. It suggested that upregulation of IHC genes was relatively easier (or more efficiently) than downregulation of OHC genes. Collectively, our data support that Tbx2 alone is not competent to fully overwrite the OHC transcriptional program and convert them into bona fide ‘IHCs’ as previously proposed ^27^. Instead, the cIHCs are likely to be, more or less, in a hybrid stage co-expressing many genes that are otherwise mutually exclusive in IHCs or OHCs in the wild type mice.

### Embryonic HC progenitors overexpressing Tbx2 develop into IHC-like cells with abnormal hair bundles

We next wondered whether Tbx2 misexpression at embryonic ages would completely transform the HC progenitors, especially those that normally would adopt OHC fate, into IHCs by briefly analyzing the *Atoh1^Cre/+^*; *Rosa26^Tbx^*^2^*^/+^* (Atoh1-Tbx2 in brief). Almost all HC progenitors and a small fraction of SC progenitors are targeted in *Atoh1^Cre/+^* ^35^. We confirmed that ectopic nucleus HA (Tbx2) expression was not detected in the HCs of control *Atoh1^Cre/+^* (Supplemental Figure 6A-A’’’), but was present in HCs of the Atoh1-Tbx2 cochleae (arrows in Supplemental Figure 6B-B’’’). Moreover, Fgf8 is a specific marker for nascent and differentiating IHCs ^36^, and GFP can faithfully represent Fgf8 expression in the knockin mouse strain *Fgf8*-P2A-3◊GFP/+ (*Fgf8^GFP/+^* in brief) ^37^. We further analyzed the control *Atoh1^Cre/+^*; *Fgf8^GFP/+^* (Atoh1-Fgf8 in short) and experimental *Atoh1^Cre/+^*; *Rosa26^Tbx2/+^*; *Fgf8^GFP/+^* (Atoh1-Tbx2-Fgf8 in short) at P2. Triple staining of GFP, tdTomato and early OHC marker Bcl11b showed that GFP (Fgf8) and Bcl11b were exclusive to IHCs and OHCs, respectively (Figure 4A-A’’’). In contrast, all tdTomato+ cells in OHC region lost Bcl11b, and started to express GFP (Fgf8) expression in Atoh1-Tbx2-Fgf8 at P2 (arrows in Figure 4B-B’’’). Likewise, those GFP+/tdTomato+/Bcl11b-cells in the OHC region were defined as cIHCs. We further analyzed *Atoh1^Cre/+^* and *Atoh1^Cre/+^*; *Rosa26^Tbx2/+^* (Atoh1-Tbx2) mice at P7. Relative to the control *Atoh1^Cre/+^*in which Prestin was exclusively expressed in OHCs and vGlut3 was specific to IHCs at P7 (Figure 4C-C’’’), in Atoh1-Tbx2 mice Prestin completely disappeared in the tdTomato+ cIHCs that instead expressed vGlut3 (arrows in Figure 4D-D’’’). The tdTomato+ IHCs seemed normal (arrowheads in Figure 4B-B’’’ and Figure 4D-D’’’), supporting that IHCs could tolerate additional Tbx2 expression.

**Figure 4.**
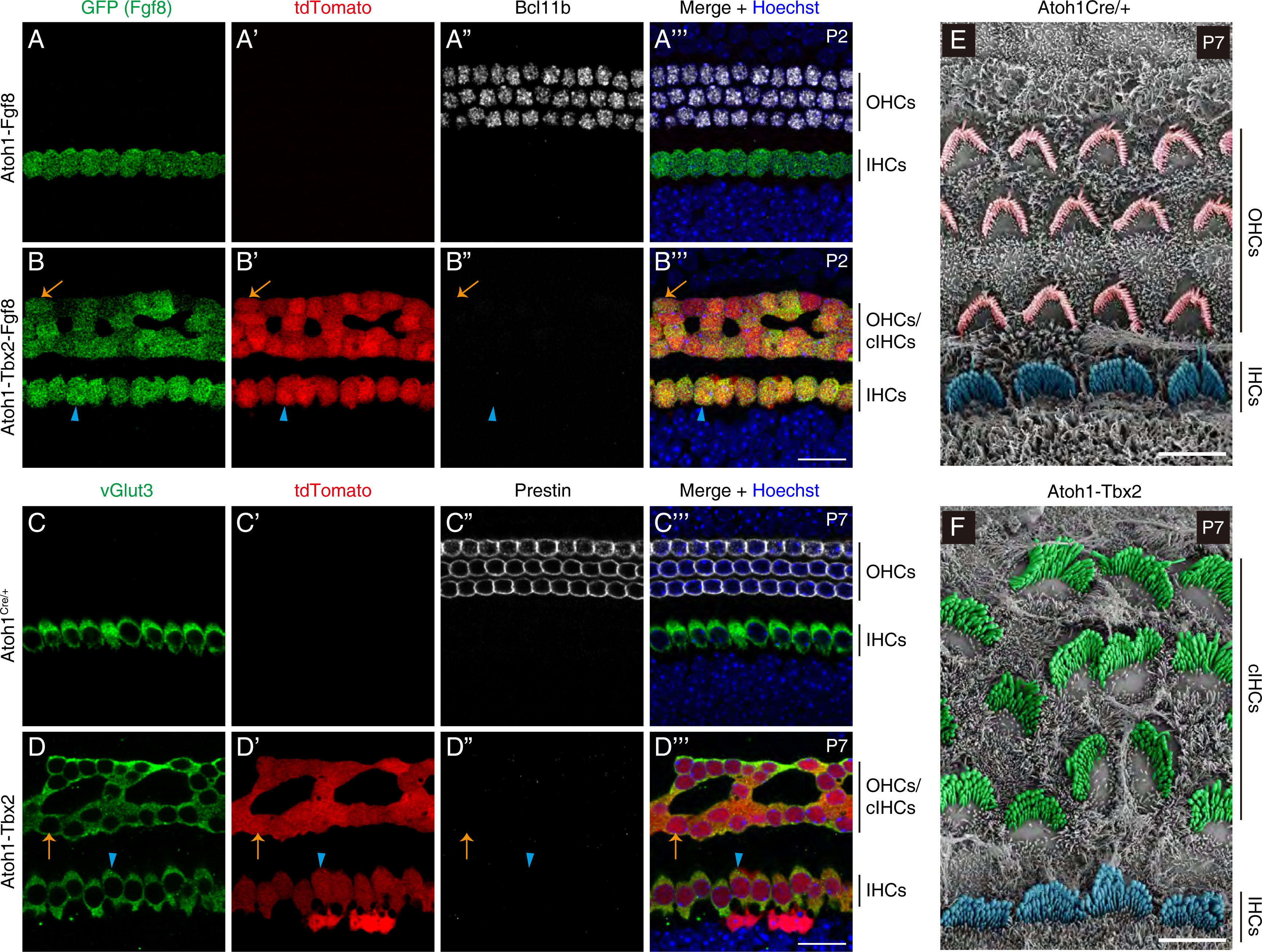
Tbx2 misexpression converts lateral cochlear progenitors into IHC-like cells with disorganized hair bundles. **(A-B’’’)** Triple staining of GFP (Fgf8), tdTomato and the OHC marker Bcl11b in cochlear samples of control Atoh1-Fgf8 (n=3, A-A’’’) and experimental group Atoh1-Tbx2-Fgf8 mice (n=3, B-B’’’) at P2. The orange arrows in (B-B’’’) point to the same cIHC expressing GFP (Fgf8), tdTomato but losing Bcl11b. The blue arrowheads mark one IHC that expresses GFP (Fgf8), tdTomato but not Bcl11b. **(C-D’’’)** Triple staining of vGlut3, tdTomato and Prestin in cochlear tissues of control *Atoh1^Cre/+^*(n=3, C-C’’’) and Atoh1-Tbx2 mice (n=3, D-D’’’) at P7. The orange arrows in (D-D’’’) label one vGlut3+/ tdTomato+/Prestin-cIHC, whereas the blue arrowheads point to one IHC. The tdTomato+ cells medial to IHCs (D-D’’’) were supporting cells that also can be targeted, but much less frequently than HCs, by the *Atoh1^Cre/+^*. **(E-F)** SEM scanning of hair bundles in *Atoh1^Cre/+^* (n=3, E) and Atoh1-Tbx2 mice (n=3, F) at P7. Pink color marks the V or W-shaped OHC hair bundles (E), green marks the IHC-like hair bundles of the cIHCs (F). The blue color marks the IHC hair bundles in both models. Scale bars: 20 μm (B’’’ and D’’’), 5 μm (E and F).

Notably, we noticed that the cIHCs alignment was not as regular as those OHCs in control mice, especially at P7 (Figure 4D-D’’’). Consistently, SEM analysis showed that, relative to OHC hair bundles (pink color in Figure 4E) in control *Atoh1^Cre/+^*mice at P7, the hair bundle orientation of the cIHCs were disorganized (green color in Figure 4F), albeit per cell they lost the V or W-shaped, instead gained the bird wing-shaped hair bundle, similar to the endogenous IHCs (blue color in Figure 4F). Collectively, our data demonstrated that, in the presence of ectopic Tbx2, the cochlear development in the lateral portion was grossly defective and we did not observe four rows of bona fide “IHCs”.

### Tbx2 misexpression in adult OHCs is able to repress Prestin and derepress vGlut3 expression

Whether Tbx2 misexpression is competent to perturb the gene expressions of adult OHCs remains completely unknown. The Slc26a5-Tbx2 mice were administered tamoxifen at P60 and P61, and analyzed at P90 or P120, in parallel with the control Slc26a5-Ai9 mice (Figure 5A). Different from control (Figure 5B-B’’’), vGlut3 was induced and Prestin was repressed in the tdTomato+ cells that were also defined as cIHCs at P90 (Figure 5C-C’’’). Both vGlut3 and Prestin expression levels were heterogeneous among the tdTomato+ cIHCs. Nevertheless, per cIHC the higher level the vGlut3 was, the lower would be the Prestin level (comparison between the orange arrow and arrowheads in Figure 5C-C’’’). Such a contrasting pattern between vGlut3 and Prestin agreed with the unsynchronized and various incomplete stages of cIHCs along the roadmap of the OHC to IHC conversion. Notably, the heterogeneity largely disappeared at P120 (Figure 5D-D’’’). All tdTomato+ cIHCs expressed high vGlut3 but faint or completely lost Prestin (orange arrowheads in Figure 5D-D’’’). As expected, the OHCs not expressing tdTomato harbored the highest level of Prestin and did not express detectable vGlut3 at P90 and P120 (blue arrows in Figure 5C-D’’’).

**Figure 5.**
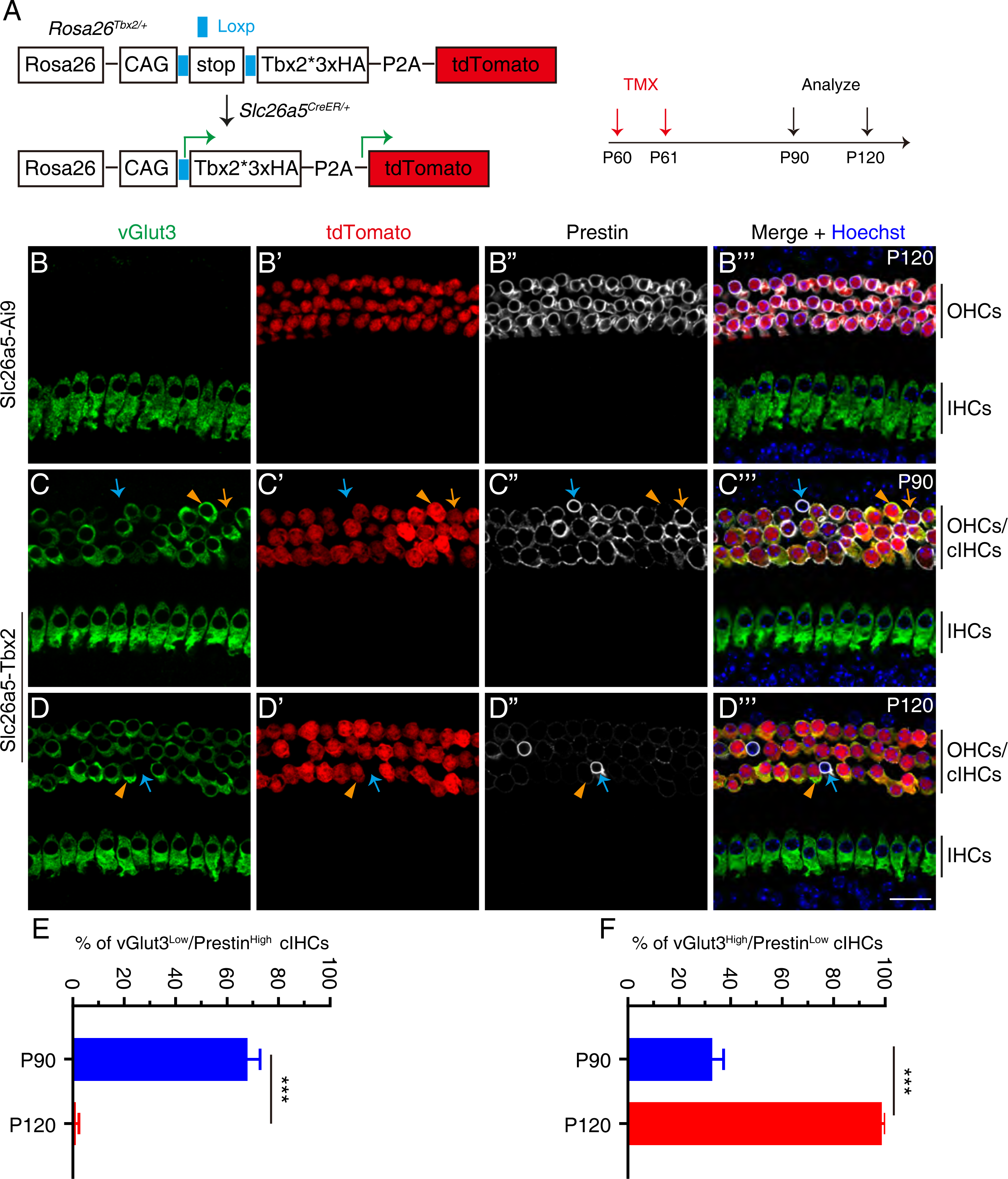
Tbx2 misexpression decreases Prestin and derepresses vGlut3 in adult OHCs. **(A)** A simple illustration of how we specifically start to induce Tbx2 expression in adult OHCs. **(B-D’’’)** Triple staining of vGlut3, tdTomato and Prestin in wholemount cochlear tissues of control Slc26a5-Ai9 at P120 (n=3, B-B’’’), Slc26a5-Tbx2 at P90 (n=3, C-C’’’) or at P120 (n=3, D-D’’’). All mice were administrated tamoxifen at P60 and P61. The orange arrows (C-C’’’) mark one tdTomato+ cIHC that expresses low level of vGlut3 but high level of Prestin, whereas orange arrowheads (C-C’’’ and D-D’’’) label the tdTomato+ cIHCs expressing high level of vGlut3 but low level of Prestin. The blue arrows (C-C’’’ and D-D’’’) point to the endogenous OHCs that do not express tdTomato (Tbx2) and vGlut3, but maintain Prestin expression level as high as the endogenous OHCs in control mice (B’’). **(E-F)** Quantification of the percentage of vGlut3^Low^/Prestin^High^ cIHCs (E) or vGlut3^High^/Prestin^Low^ cIHCs (F). Data are presented as Mean ± Sem. Student’s *t* test was used for statistical analysis. ***p<0.001. Scale bar: 20 μm (D’’’).

To simplify the quantification, here we roughly divided cIHCs into two categories, vGlut3^Low^/Prestin^High^ (orange arrows in Figure 5C-C’’’) and vGlut3^High^/Prestin^Low^ (orange arrowheads in Figure 5C-D’’’). Here, the ‘Low’ included both faint or undetectable protein expression. Among all tdTomato+ cIHCs, 67.95% ± 4.87% (n=3, mouse number) were vGlut3^High^/Prestin^Low^ and 32.77% ± 4.48% (n=3, mouse number) were vGlut3^Low^/Prestin^High^ at P90, whereas almost all (98.75% ± 1.25%, n=3, mouse number) were vGlut3^High^/Prestin^Low^ at P120 (Figure 5E and F). In sum, we concluded that Tbx2 misexpression alone was sufficient to, at least to some degree, perturbate the gene expressions in adult OHCs.

### Construction of a new mouse strain in which Tbx2 and Ikzf2 can be simultaneously overexpressed

The molecular mechanism underlying how Tbx2 destabilizes the OHC fate remains completely unknown. Both *Ikzf2* mRNA and protein were repressed in the cIHCs (Supplemental Figure 3H-H’’ and Supplemental Figure 5B). Because Ikzf2 is necessary in consolidating the OHC fate ^22^, we hypothesized that repression of Ikzf2 by Tbx2 was one of the key contributors for the cIHC production. We tested this hypothesis by determining whether Ikzf2 restoration could mitigate the effects of Tbx2 on OHCs.

In order to guarantee that Ikzf2 would be simultaneously restored in all OHCs expressing Tbx2, we constructed a new *Rosa26*-Loxp-stop-Loxp-Tbx2*3◊HA-P2A-Ikzf2*3 ◊V5-T2A-EGFP/+ (*Rosa26^Tbx2-Ikzf2/+^* in brief) mouse line (Figure 6A and Supplemental Figure 7A-C). No random insertion of the targeting vector happened in mouse genome, as revealed by the Southern blot assay (Supplemental Figure 7D). The wild type (WT) and heterozygous knockin (KI) *Rosa26^Tbx2-Ikzf2/+^* were readily to be identified by tail DNA PCR (Supplemental Figure 7E). Notably, after cre-mediated recombination, OHCs expressing Tbx2 and Ikzf2 would be EGFP+, as to be confirmed below.

**Figure 6.**
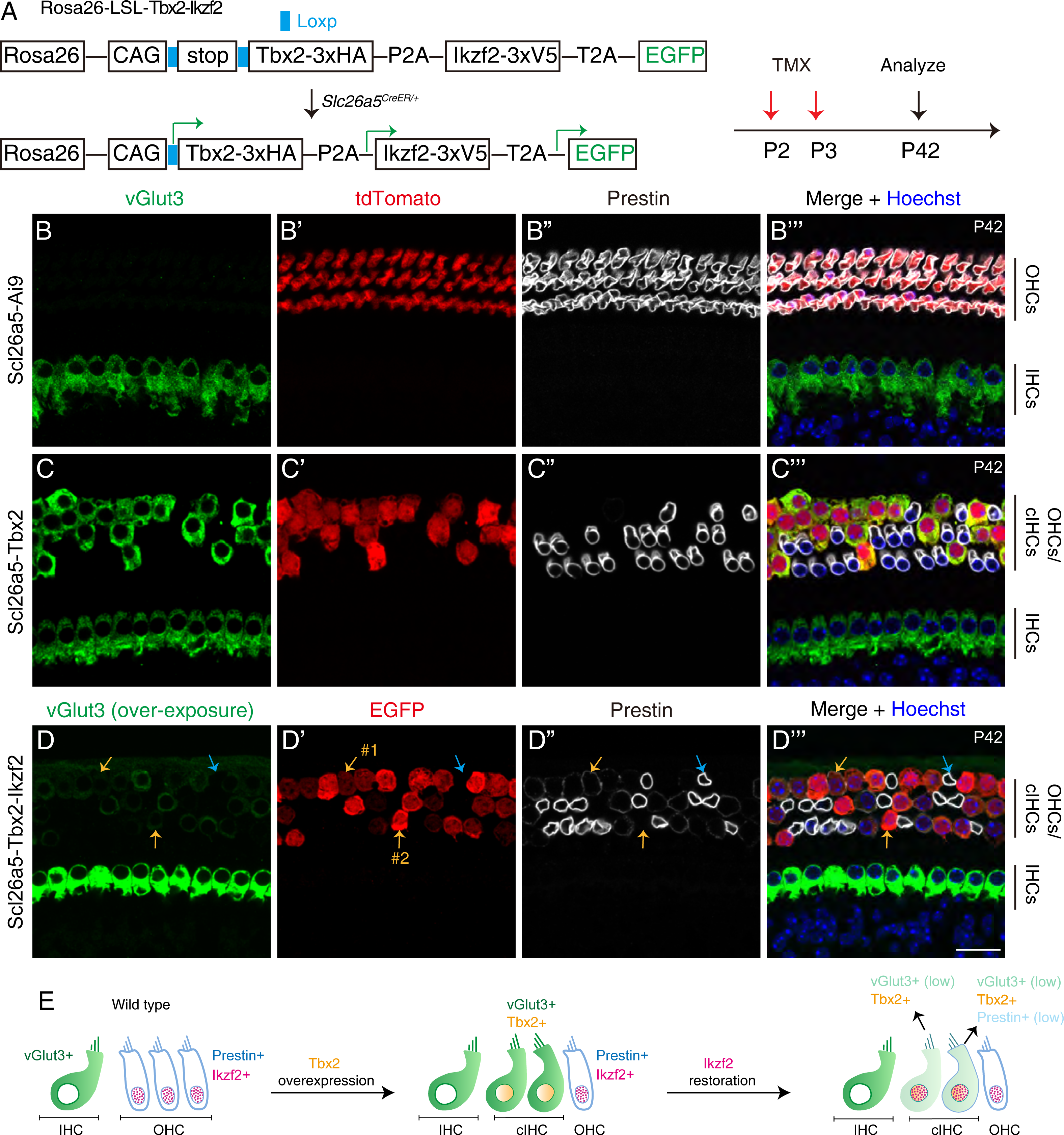
Ikzf2 antagonizes the effects of Tbx2 on reprogramming OHCs into IHC-like cells or cIHCs. **(A)** A simple cartoon to show how we simultaneously induce Tbx2, Ikzf2 and EGFP expression in OHCs at P2 and P3. **(B-D’’’)** Triple staining of vGlut3, tdTomato (or EGFP) and Prestin in control Slc26a5-Ai9 (B-B’’’, n=3) and Slc26a5-Tbx2 (C-C’’’, n=3), as well as the Slc26a5-Tbx2-Ikzf2 (D-D’’’, n=3) mice at P42. The vGlut3 level in the cIHCs in Slc26a5-Tbx2-Ikzf2 mice is weak or undetectable, and the vGlut3 channel must be overexposed in IHCs in order to visualize the vGlut3 in the cIHCs. The orange arrows (#1, D-D’’’) mark one EGFP+ cIHC that expresses low level of Prestin. The orange arrows (#2, D-D’’’) mark one EGFP+ cIHC in which Prestin is undetectable. Blue arrows (D-D’’’) label one endogenous OHC that does not express EGFP (Ikzf2 and Tbx2), and maintain Prestin expression as high as the OHCs in control mice (B-B’’’). **(E)** A summarized model to highlight that forced Ikzf2 expression in the cIHCs mitigates the abnormal vGlut3 expression, as well as partially restores Prestin expression in the cIHCs. Scale bar: 20 μm (D’’’).

### Ikzf2 restoration antagonizes the effect of Tbx2 on converting OHCs into IHC-like cells

By analyzing the *Slc26a5^CreER/+^*; *Rosa26^Tbx2-Ikzf2/+^* (Slc26a5-Tbx2-Ikzf2 in brief) at P42, we confirmed that all EGFP+ cells in the OHC region expressed the nucleus HA (Tbx2) and V5 (Ikzf2) (orange arrows in Supplemental Figure 7F-F’’’), whereas the EGFP-OHCs did not express nucleus HA (Tbx2) and V5 (Ikzf2) (blue arrows in Supplemental Figure 7F-F’’’). Notably, the membrane HA signal was contributed by the HA-tagged Prestin in *Slc26a5^CreER/+^* mice ^29^. We continued defining those EGFP+ cells in Slc26a5-Tbx2-Ikzf2 model as cIHCs, although the cIHCs should be different from the cIHCs described in the Slc26a5-Tbx2 model above.

Next, triple straining of vGlut3, tdTomato (or EGFP) and Prestin were applied to three models: 1) Slc26a5-Ai9 (Figure 6B-B’’’); 2) Slc26a5-Tbx2 (Figure 6C-C’’’); and 3) Slc26a5-Tbx2-Ikzf2 (Figure 6D-D’’’). The first two models had been described above and here only served as controls. Briefly, two lines of evidence supported that Ikzf2 markedly, albeit not completely, antagonized the effects of Tbx2 on the neonatal OHCs. First, vGlut3 expression levels in all EGFP+ cIHCs were drastically decreased or barely detected, and the vGlut3 channel in IHCs had to be overexposed to visualize the weak vGlut3 (Figure 6D-D’’’). Second, Prestin expression was restored in 79.01% ± 5.59% of the EGFP+ cIHCs (orange arrows in Figure 6D-D’’’, #1), but the expression level was much weaker than that of the nearby EGFP-endogenous OHCs (blue arrows in Figure 6D-D’’’). However, Prestin was undetectable in the remaining 20.99% ± 5.59% of the EGFP+ cIHCs (orange arrows in Figure 6D-D’’’, #2). Collectively, relative to the cIHCs misexpressing Tbx2 only, Ikzf2 restoration can partially reverse the cIHCs into a status closer to OHCs, with repressing vGlut3 and increasing Prestin (Figure 6E). Thus, repression of Ikzf2 by Tbx2 misexpression is one of the key transcriptional cascades involved in destabilizing the OHC fate.

## Discussion

### Specification of cochlear progenitors into IHCs or OHCs

Figuring out the molecular mechanisms underlying how cochlear progenitors develop into IHCs or OHCs is of importance before establishing an efficient *in vivo* approach to regenerate HCs after trauma. It has been granted for a long time that there are pan-HC progenitors from which IHCs and OHCs are born ^9,10^. Those pan-HC progenitors in general express higher Atoh1 than those that otherwise would become SCs ^18^. However, recent transcriptomic and genetic evidence support the second proposal that IHCs share the same progenitors with SCs on the medial side, and that OHCs and lateral SCs derive from the same progenitors ^38,39^. The second proposal is further supported by the expression pattern of Tbx2 which is broadly expressed in cochlear cells before E13.5, but is gradually diminished, following a basal to apical gradient, in the lateral cochlear progenitor cells ^26–28^. In contrast, Tbx2 is persistently expressed in medial progenitors that would differentiate into IHCs or SCs including inner border cells (IBCs) and inner phalangeal cells (IPhs) ^26,28^. Our current study, together with previous studies ^26–28^, support that Tbx2 blocks the cell fates of the lateral auditory epithelium, including OHCs, and nearby two SC subtypes, Pillar and Deiters’ cells.

How does Tbx2 promote medial progenitors to differentiate into IHCs? We proposed that Tbx2+ progenitors further expressing high level of Atoh1 protein would differentiate into IHCs. Our proposal is supported by the report that Tbx2 and Atoh1 together successfully reprogram IBCs and IPhs into vGlut3+ IHCs ^26,40^. On the other hand, how do the Tbx2-lateral progenitors become OHCs? Because Tbx2 diminishment occurs earlier than the onset of Insm1 in the nascent OHCs ^21,26^, we suspected that absence of Tbx2 in lateral progenitors should create a permissive environment for Insm1 expression. Insm1 primarily acts as a transcriptional repressor and blocks IHC gene expression in nascent OHCs ^20^. Such a proposal is further supported by that Insm1 is derepressed in the *Tbx2^-/-^* IHCs ^27^. Thus, cochlear progenitors that express both Insm1 and high level of Atoh1 protein become OHCs.

### Molecular mechanism underlying OHC differentiation and fate maintenance at adult ages

Insm1 should be only involved in the early stage of OHC development, because it is transiently expressed in the nascent OHCs in a basal to apical gradient and becomes undetectable by P2 ^21,41^. Then, how is OHC differentiation regulated at ages after P2? Although much remains unclear, Ikzf2 clearly plays critical roles in it. Ikzf2 is turned on in OHCs at ∼P4 and maintained afterwards ^22,26^. The defective *Ikzf2^cello/cello^*OHCs are dysfunctional and degenerate, resulting in early-onset sensorineural hearing loss ^22^. Notably, we recently report that Ikzf2 is positively but indirectly regulated by Insm1, and restoration of Ikzf2 partly alleviates the abnormalities of the *Insm1^-/-^* OHCs ^21^. Thus, the existence of the transcriptional signaling cascade, from Tbx2 to Insm1 and from Insm1 to Ikzf2, might explain why Ikzf2 restoration can partially mitigate the effects of Tbx2 misexpression on OHCs, as revealed by our analyses (Figure 6). Nonetheless, it remains unknown whether Insm1 restoration also can antagonize Tbx2 misexpression in OHCs. It is also interesting to know whether dual restoration of Insm1 and Ikzf2 can block the effects of Tbx2 on OHCs more effectively than Insm1 or Ikzf2 alone.

It is known that neonatal OHCs transdifferentiate into IHC-like cells upon forced Tbx2 expression ^27,28^. Our study extends to show that Tbx2 is competent to destabilize the adult OHC fate, with upregulating vGlut3 and repressing Prestin expression as the readouts. Given reprogramming adult fully differentiated cells is much more difficult than doing young and immature counterparts, Tbx2 should be regarded as powerful reprogramming factor. Due to lack of *Ikzf2* conditional loss-of-function study, the precise roles of Ikzf2 in adult OHCs remain to be determined. It is possible that Ikzf2 continues maintaining the OHC fate at adult ages, and repression of Ikzf2 by Tbx2 in adult OHCs should contribute to their perturbed gene expressions.

### Degeneration of hair cells in the presence of ectopic Tbx2 expression

Previous *Tbx2* gain-of-function studies focus on whether Tbx2 is competent to reprogram neonatal OHCs into IHCs or IHC-like cells ^27,28^. The long-term effect of Tbx2 on OHCs are not addressed. Our study showed that the cIHCs started to degenerate after P14 in a basal to apical gradient (Supplemental Figure 2). Why did the cIHCs undergo cell death? We provided two interpretations: 1) The degeneration is caused by Prestin repression in the cIHCs. It is well known that *Prestin^-/-^* OHCs have normal development at early ages, but gradually degenerate at adult ages ^4^; 2) Loss of Ikzf2 function leads to OHC degeneration in the *Ikzf2^cello/cello^* mice ^22^. Thus, it is possible that the repression of Ikzf2 directly contributes to the degeneration of cIHCs. Notably, the precise mechanisms underlying degeneration of the *Prestin^-/-^* and *Ikzf2^cello/cello^* OHCs remains poorly understood. Moreover, the IHCs and OHCs not overexpressing Tbx2 also degenerated after P42, which should be a second effect due to the cell death of the cIHCs.

### Tbx2 is a powerful transcription factor but should not be able to fully reprogram cochlear progenitors or OHCs into IHCs

Three lines of evidence support that Tbx2 is a more upstream and powerful transcription factor than Insm1 and Ikzf2 during HC development. First, while only ∼50% of *Insm1^-/-^* OHCs become IHC-like cells ^20,21^, almost all *Tbx2^-/-^* IHCs become OHC-like cells ^26^. Second, *Tbx2* is epistatic to *Insm1* and *Tbx2^-/-^; Insm1^-/-^* IHCs transdifferentiate into OHC-like cells ^27^; Third, restoration of Ikzf2 cannot fully restore the OHC features in the Tbx2+ cIHCs (Figure 6). Recently, the mechanism of how Tbx2 binds specific DNA sequences, and its association co-factors are reported in other cell contexts ^42^. However, such a dataset of Tbx2 in cochlear cells is not available yet.

Nonetheless, different from the previous report ^27^, five of our observations below support that Tbx2 cannot fully reprogram other cochlear cell types into bona fide IHCs: 1) our full-length single cell transcriptomic assay clearly showed that only 38.6% of OHC genes were significantly repressed in the cIHCs (Figure 3) and manifest global gene expression difference existed between cIHCs and IHCs (Supplemental Figure 5D); 2) the cIHCs harbored degenerated or lost hair bundles (Figure 2E and F); 3) the overall cochlear development was disruptive when Tbx2 was overexpressed in cochlear sensory progenitors, the cIHCs displayed disorganized hair bundles (Figure 4); 4) the number of the ribbon synapses in the cIHCs was much less than in the endogenous IHCs (Supplemental Figure 3); 5) the total cell surface area of cIHCs was smaller than that of OHCs (Figure 2), thus should also be smaller than that of IHCs. Collectively, we speculated that, although Tbx2 is one of the key IHC fate determinators, additional genes are needed to fully reprogram OHCs or HC progenitors into IHCs, especially into IHCs harboring the well-organized IHC-like hair bundles.

## MATERIALS AND METHODS

### Mice

The *Slc26a5^CreER/+^* is kindly provided by Dr. Jian Zuo (Creighton University, USA)^29^. The *Atoh1^Cre/+^* is kindly provided by Dr. Lin Gan (Augusta University, USA) ^35^. The details of *Fgf8^GFP/+^* strain are described in our previous report ^37^. Tamoxifen (TMX) (Cat# T5648, Sigma-Aldrich) was dissolved in corn oil (Cat# C8267, Sigma-Aldrich) and administrated into mice, with the dose of 3 mg/40 g (body weight) at P2/P3 or 9 mg/40 g (body weight) P60/P61. All mice were bred and raised in a SPF-level animal room and all animal procedures were performed according to the guidelines (NA-032-2022) of the IACUC of the Institute of Neuroscience (ION), Center for Excellence in Brain Science and Intelligence Technology, Chinese Academy of Sciences.

### Generation of Rosa26^Tbx^^2/+^ and Rosa26^Tbx^^2-Ikzf^^2/+^ strains

The *Tbx2* conditional overexpression mouse strain *Rosa26*-Loxp-stop-Loxp-Tbx2*3 ◊ HA-P2A-tdTomato (*Rosa26^Tbx^*^2^*^/+^*) was established by the homologous recombination mediated by CRISPR/Cas9 in one-cell stage mouse zygotes into which *Cas9* mRNA, targeting vector and *Rosa26* sgRNA were coinjected (Supplemental Figure 1). The *Rosa26* sgRNA is: *5ʹ-ACTCCAGTCTTTCTAGAAGA-3ʹ*. The injected zygotes were subsequently transplanted into pseudopregnant female mice that would give birth to the founder 0 (F0) mice. F0 mice were screened by tail PCR and those with potential correct gene targeting were bred with wild type mice to produce the germ line stable F1 mice. Southern blot assay was performed to show that there was no random insertion of the targeting vector in the F1 mouse genome (Supplemental Figure 1D). The detailed protocol of Southern blot is reported in our previous report ^43^. Likewise, the same procedure was used to generate the *Rosa26*-Loxp-stop-Loxp-Tbx2*3 ◊ HA-P2A-Ikzf2*3 ◊ V5-T2A-EGFP/+ (*Rosa26^Tbx^*^2^*^-Ikzf^*^2^*^/+^*) except that the targeting vector was different from *Rosa26^Tbx2/+^*.

Same mouse genotyping procedures were applied to both *Rosa26^Tbx2/+^* and *Rosa26^Tbx2-Ikzf2/+^*. We concurrently used three primers (F1, R1, and R2 in Supplemental Figures 1 and 7) to distinguish the KI and WT alleles. The primer sequences were: Forward primer 1 (F1): *5ʹ-AGTCGCTCTGAGTTGTTATCAG-3ʹ*; Reverse primer 1 (R1): *5ʹ-TGAGCATGTCTTTAATCTACCTCGATG-3ʹ*; Reverse primer 2 (R2): *5ʹ-AGTCCCTATTGGCGTTACTATGG-3ʹ.* The following PCR protocol was used: 95°C for 3 min, followed by 30 cycles of 95°C for 30 s, 60°C for 30 s, and 72°C for 40 s, and then a final extension at 72°C for 10 min. Whereas the paired primers F1 and R1 produced a 469 bp fragment from WT allele, the paired primers F1 and R2 yielded a 412 bp fragment from the KI allele.

### Sample processing and immunofluorescence assessment

The detailed protocol of inner ear tissue processing is reported in our previous studies ^44^. Briefly, after the inner ear tissues were dissected out, they were fixed in fresh 4% paraformaldehyde (PFA) overnight at 4°C. On the following morning, they were washed thrice for 10 min each in 1× PBS, followed by decalcification in 120 mM EDTA at room temperature.

The primary antibodies used in this study included anti-HA (rat, 1:200; 11867423001, Roche), anti-Prestin (goat, 1:1000; sc-22692, Santa Cruz), anti-vGlut3 (rabbit, 1:500; 135203, Synaptic Systems), anti-GFP (chicken, 1:500; ab13970, Abcam), anti-Bcl11b (rat, 1:500; ab18465, Abcam), anti-Otoferlin (mouse, 1:500; ab53233, Abcam), anti-Slc7a14 (rabbit, 1:500; HPA045929, Sigma-Aldrich), anti-Ctbp2 (mouse, 1:200; 612044, BD Biosciences), anti-V5 (mouse, 1:500; MCA1360, Bio-Rad). Nuclei was counterstained by Hoechst33342 solution (1:1000; 62249, Thermo Scientific). Prolong gold antifade medium (P36930; Thermo Scientific) was used for sample mounting. NiE-A1 Plus, TiE-A1 Plus and Nikon C2 (Nikon, Japan) confocal microscopes were used for image capture.

### Cell number, ribbon synapse and nuclei diameter quantification and measurement

Each cochlea was divided into three portions, basal, middle and apical turn with equal length after measuring its total length scanned under the 10× lens of the confocal microscope. Numbers of the surviving cells (OHCs or cIHCs) in control (Slc26a5-Ai9) and experimental group (Slc26a5-Tbx2) were counted at basal turns at P42, at basal/middle turns at P120, respectively (Supplemental Figure 2). The numbers of the Ctbp2+ puncta were used to calculate the ribbon synapse. The mosaic distribution of the endogenous OHCs and the cIHCs in the Slc26a5-Tbx2 mice permitted us to compare the numbers of the Ctbp2+ puncta in cIHCs, OHCs as well as the IHCs in the same region, minimizing the contribution of cochlear region difference to the varied number of the Ctbp2+ puncta (Supplemental Figure 3). The same approach was applied to measure the nucleus diameter of cIHCs, OHCs and IHCs in Slc26a5-Tbx2 mice (Supplemental Figure 3).

To quantify the percentages of vGlut3+/Prestin^Low^ or vGlut3+/Prestin^-^ cIHCs in Slc26a5-Tbx2 mice at P14, P42, P76 and P120 (Figure 1), we chose the apical turn because none or very limited cIHCs/OHCs were lost. The percentage was calculated by normalizing the number of vGlut3+/Prestin^Low^ or vGlut3+/Prestin^-^ cIHCs to the total number of the Tdtomato+ cells in the OHC region, which was also the sum of vGlut3+/Prestin^Low^ or vGlut3+/Prestin^-^ cIHCs. The same method was applied to quantify the percentages of vGlut3^Low^/Prestin^High^ or vGlut3^High^ /Prestin^Low^ cIHCs in the Slc26a5-Tbx2 mice in which Tbx2 was induced at P60/P61 (Figure 5).

### Bioinformatics analysis

FASTQ files of smart-seq data were aligned to the house mouse reference genome (GRCm38) using Hisat2 (v2.2.1) ^45^. In order to estimate the expression level of each gene, raw count was calculated by featureCounts (v2.0.1) ^46^, and normalized value TPM (transcript per million) was calculated by StringTie (v2.2.1) ^47^. Differentially expressed genes (DEGs) between P42_cIHCs and P30_WT IHCs/OHCs were identified by DESeq2 (v1.34.0) ^48^, with the criteria: absolute value of FC > 4, p < 0.05. Genes in lists of supplemental tables 1 and 2 were ranked in order of averaged TPM.

Moreover, R package Seurat (v4.0.6) was utilized for other downstream analysis^49^. Principal component analysis (PCA) was carried out by “RunPCA” and clustering was performed by “FindNeighbors” and “FindClusters”. The dimensional reduction, uniform manifold approximation and projection (UMAP), was carried out by “RunUMAP”. All of single cell transcriptomic data *de novo* generated in this study could be accessed with the number: GSE233559.

### ABR measurement and SEM analysis

After P42 WT and Slc26a5-Tbx2 mice were anesthetized, they were transferred into a soundproof chamber, and stimulated by a range of sound intensities from 90 dB SPL to 0 dB SPL at 4, 5.6, 8, 11.3, 16, 22.6, 32 and 45 kHz. The responses were recorded by BioSigRZ software (Tucker-Davis Technologies, Inc., Alachua, FL, USA). The details of the procedure are described in our previous report ^43^.

The cochlear samples subjected to SEM analysis were processed according to the same protocols reported in our previous study ^23^. Briefly, the inner ears were fixed by 2.5% glutaraldehyde (G5882, Sigma-Aldrich), followed by decalcification in 50mM EDTA in 1× PBS. We next dissected the cochlea into three turns, with the HCs being exposed and post-fixed by 1% osmium tetroxide (18451, Ted Pella). Finally, cochlear samples were treated in a turbomolecular pumped coater (Model: Q150T ES, Quorum) and scanned in a field-emission SEM instrument (Model: GeminiSEM 300, ZEISS).

### Patch-clamp of OHCs and cIHCs

The whole-cell patch clamp recording was conducted with the Axon 200B and interfaced by an Axon Digidata 1440A using jClamp software (www.scisoftco.com) at room temperature. Individual first-row OHCs/cIHCs at 8-11 kHz region were recorded. The extracellular/intracellular solutions and detailed protocol of Patch-clamp are reported in our previous study ^5^.

NLC was measured using a continuous high-resolution two-sine stimulus protocol which involved superimposing 10 mV peak amplitude sine waves of 390.6 and 781.2 Hz onto a 300 ms voltage ramp spanning from +150 to -150 mV. The capacitance data were fitted to the first derivative of a two-state Boltzmann function.

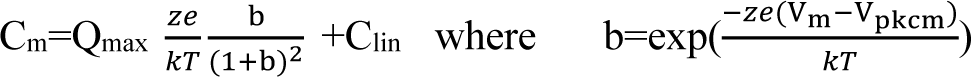

Qmax is maximum nonlinear charge moved, Vpkcm or Vh is voltage at peak capacitance, Vm is membrane potential, z represents the valence, Clin corresponds to the linear membrane capacitance, e is electron charge, k represents Boltzmann’s constant, and T is the absolute temperature. Statistical analysis was performed using one-way ANOVA with GraphPad Prism8 software.

## ACKNOWLEDGMENTS

We thank Dr. Qian Hu of the Optical Imaging Facility of the Institute of Neuroscience (ION) for support with image analysis; Drs. Xu Wang and Yu Kong from the Electronic Microscope Facility of ION for SEM assistance; and Ms. Qian Liu from the Department of Embryology of ION animal center for helping us in transplanting zygotes into pseudopregnant female mice. We thank Dr. Jian Zuo (Creighton University, USA) for kindly providing the *Slc26a5^CreER/+^* and Dr. Lin Gan (Augusta University, USA) for providing the *Atoh1^Cre/+^* strain. This study was funded by the National Key R&D Program of China (2022ZD0207000, 2020YFE0205900 and 2021YFA1101804), the National Natural Science Foundation of China (82101217 and 82000985), Strategic Priority Research Program of Chinese Academy of Science (XDB32060100), Shanghai Municipal Science and Technology Major Project (2018SHZDZX05), and Innovative Research Team of High-Level Local Universities in Shanghai (SSMU-ZLCX20180601).

## Supplemental Figure Legends

**Supplemental Figure 1.**
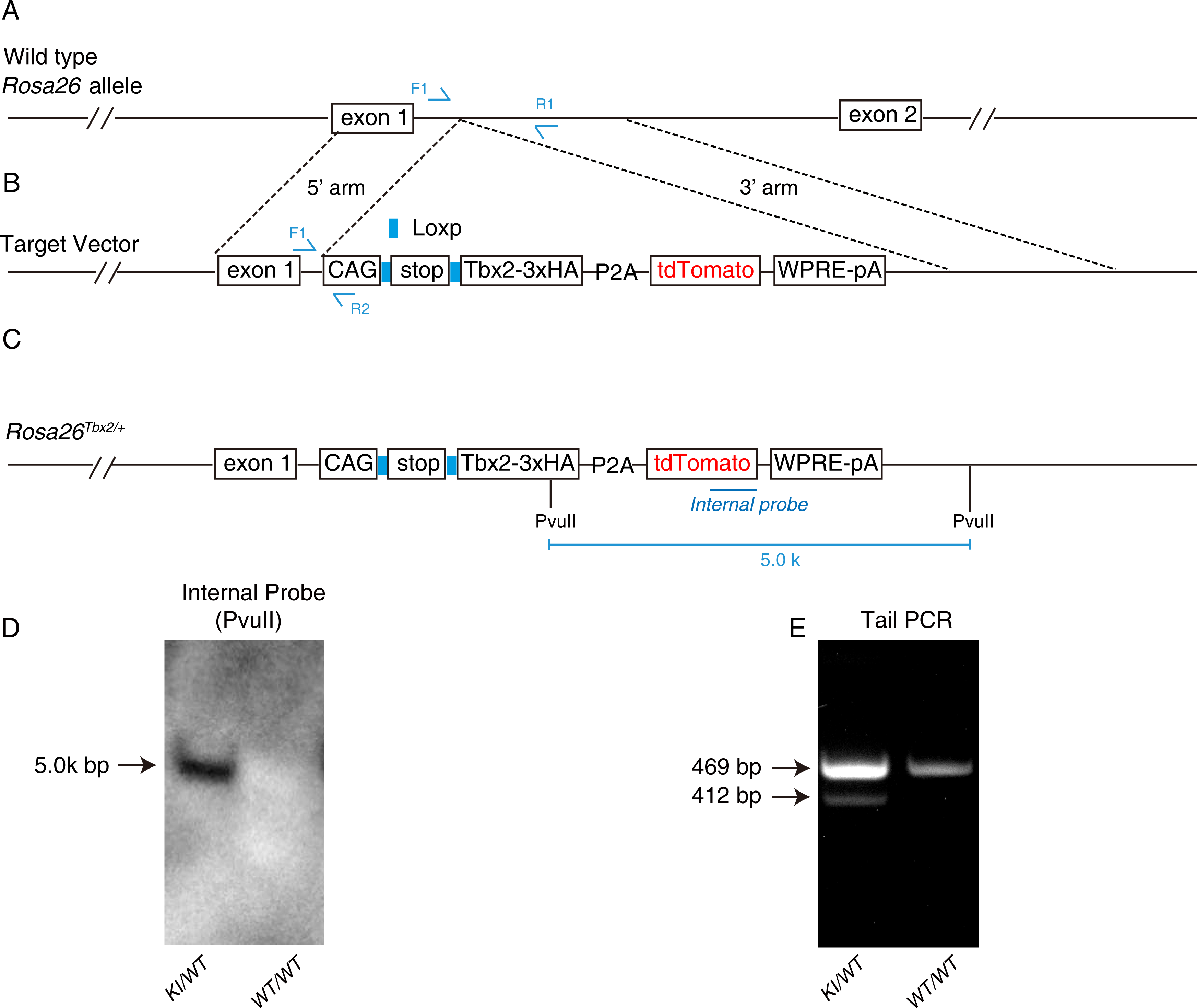
Construction of the *Rosa26*-Loxp-stop-Loxp-Tbx2*3 ◊ HA-P2A-tdTomato (*Rosa26^Tbx^*^2^*^/+^*). **(A-C)** The wild type *Rosa26* locus (A) is recombined by a targeting vector (B) with 5’ and 3’ homologous arms, resulting a post-targeted *Rosa26^Tbx^*^2^*^/+^* allele (C). Notably, Tbx2 and tdTomato expressions are tightly paired. **(D)** Southern blot assay with the internal probe. A 5k bp band is detected in the heterozygous (KI/WT), but not in the wild type (WT/WT) mice. **(E)** One exampled gel image of tail PCR of heterozygous and wild type mice.

**Supplemental Figure 2.**
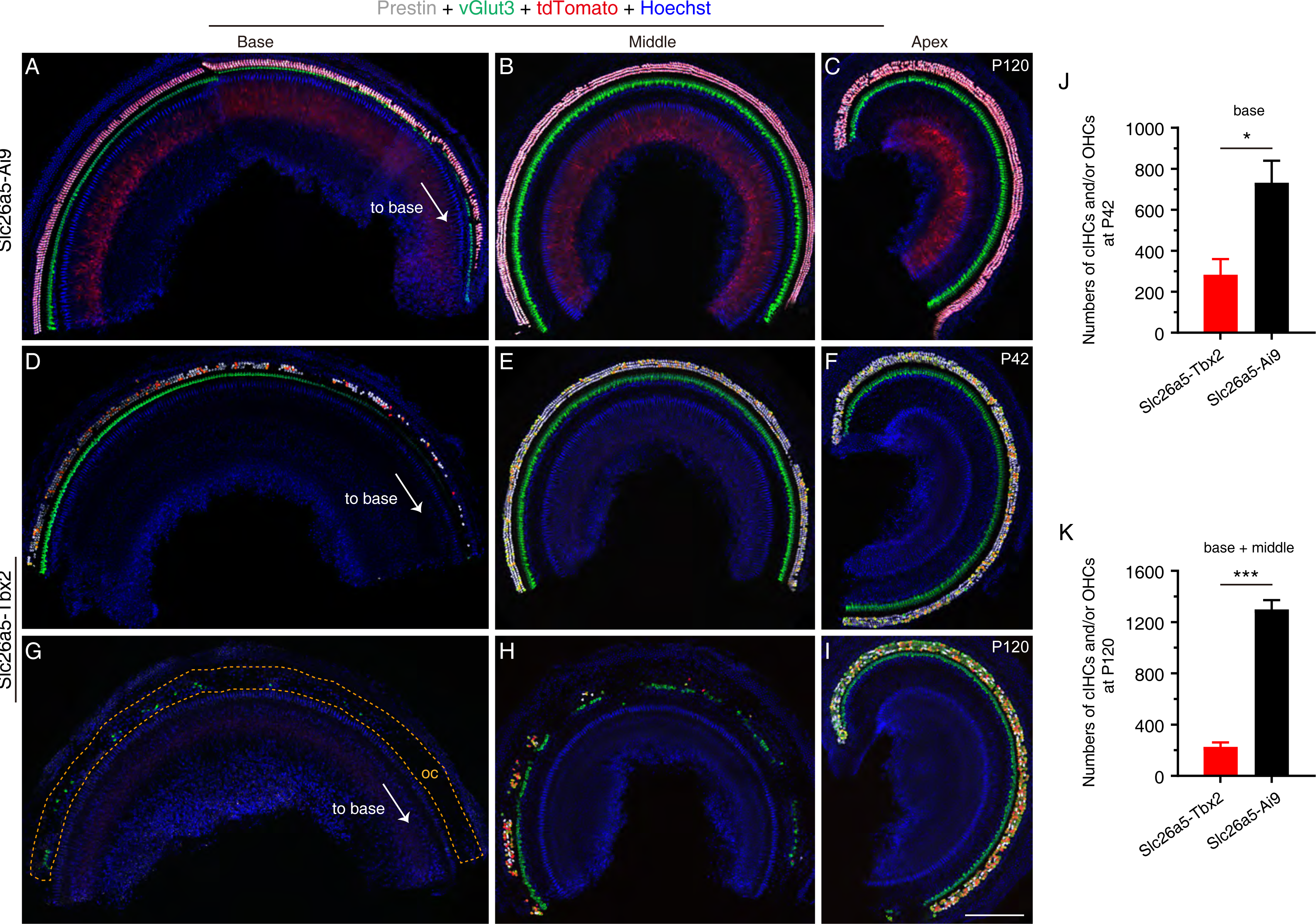
HCs are degenerated in Slc26a5-Tbx2 mice. **(A-I)** Triple staining of Prestin, vGlut3 and tdTomato in control Slc26a5-Ai9 mice (n=3) at P120 (A-C), and Slc26a5-Tbx2 mice at two different ages, P42 (D-F, n=3) and P120 (G-I, n=3). Mild HC degeneration exists in control mice, but only in the most basal portion (A). In contrast, the cIHCs or the OHCs in Slc26a5-Tbx2 start degeneration in a basal to apical gradient. The degeneration primarily happens in the basal turn (D) at P42, however, migrated to middle turn (H) at P120, but the HC degeneration in apical turn is mild at P120 (I). IHCs are normal at P42 (D-F) but are severely lost in basal (G) and middle (H), but not apical turn (I) at P120. The Tdtomato signal medial to IHCs in the control mice (A-C) was background, which was manifest in mice over 2 months old and also present in Ai9/+ mice at P120. **(J-K)** Quantification of the remaining cIHCs and OHCs in basal turns of control (black) and Slc26a5-Tbx2 (red) mice at P42 (J), as well as in basal and middle turns at P120 (K). Data are presented as Mean ± Sem. Student’s *t* test is used for statistical analysis. * p<0.05; ***p<0.001. Scale bar: 200 μm (I).

**Supplemental Figure 3.**
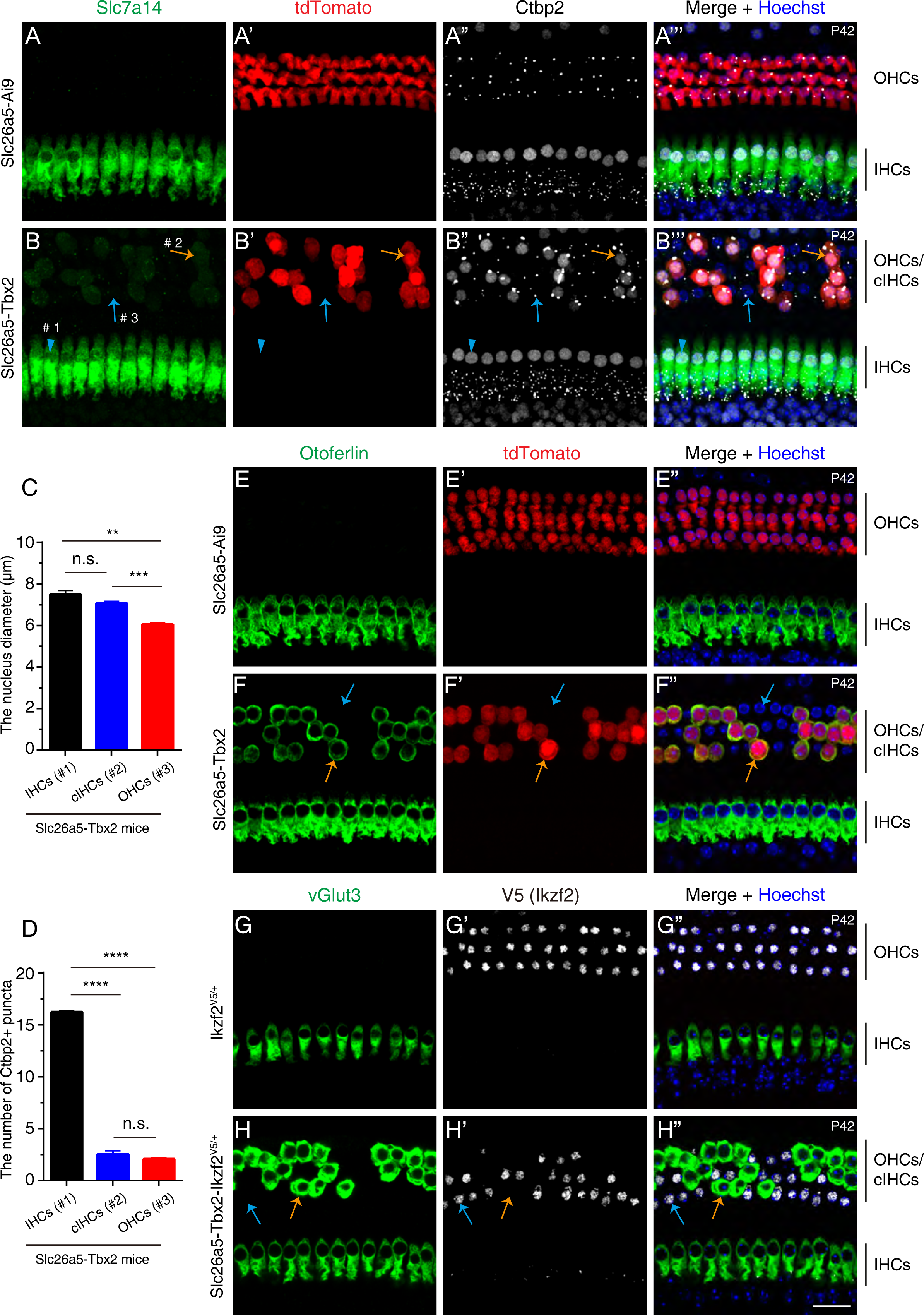
The cIHCs gain other IHC features and lose Ikzf2 protein expression. **(A-B’’’)** Triple staining of Slc7a14, tdTomato and Ctbp2 in control Slc26a5-Ai9 (A-A’’’, n=3) and Slc26a5-Tbx2 (B-B’’’, n=3) mice at P42. Blue arrowheads (#1, B-B’’’) point to one IHC, orange arrows (#2, B-B’’’) mark one tdTomato+/Slc7a14+ cIHC that expresses Ctbp2 in nuclei, and the blue arrows (#3, B-B’’’) label one endogenous OHC that does not express tdTomato (Tbx2), Slc7a14, and Ctbp2 in nuclei. **(C-D)** Among the three kinds of cell types #1, #2, and #3 in (B-B’’’), their nucleus diameters (C) and numbers of Ctbp2+ puncta in them (D) are compared. Data are presented as Mean ± Sem. Student’s *t* test is used for statistical analysis. **p<0.01; ***p<0.001; ****p<0.0001. **(E-F’’)** Co-staining of Otoferlin and tdTomato in Slc26a5-Ai9 (E-E’’, n=3) and Slc26a5-Tbx2 (F-F’’, n=3) mice at P42. Orange arrows (F-F’’) mark one tdTomato+/Otoferlin+ cIHC, whereas blue arrows (F-F’’) label one endogenous OHC that expresses neither tdTomato nor Otoferlin. **(G-H’’)** Co-staining of vGlut3 and V5 (Ikzf2) in the control *Ikzf2^V^*^5^*^/+^* mice (G-G’’, n=3) and Slc26a5-Tbx2-Ikzf2^V5/+^ mice (H-H’’, n=3) at P42. Orange arrows (H-H’’) mark one cIHC that expresses vGlut3 but loses the V5 (Ikzf2) expression, whereas blue arrows (H-H’’) point to one endogenous OHC that keeps Ikzf2 expression and does not express vGlut3. Scale bar: 20 μm (H’’).

**Supplemental Figure 4.**
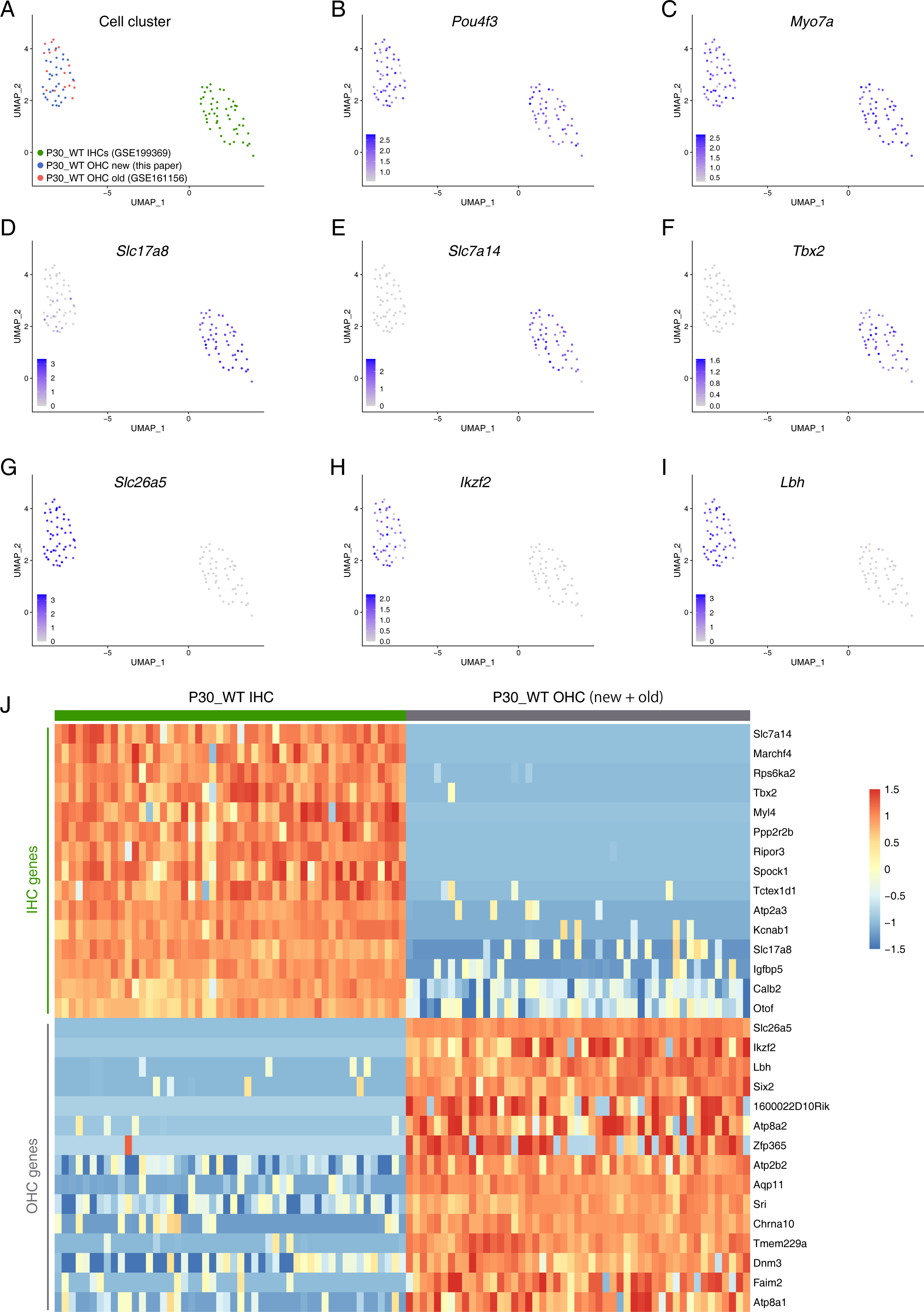
Transcriptomic difference between wild type OHCs and IHCs at P30. **(A)** Clustering of cell population that are mixed by our newly picked P30_WT OHCs new (blue, 32, cell number), and previously reported P30_WT OHCs old (red, 17, cell number) and P30_WT IHCs (green, 50, cell number). P30_WT OHCs new and P30_WT OHCs old are clustered together, and divergent from P30_WT IHCs. **(B-I)** In addition to the pan HC marker *Pou4f3* (B) and *Myo7a* (C), IHC markers *Slc17a8* (D), *Slc7a14* (E) and *Tbx2* (F), as well as OHC marks *Slc26a5* (G), *Ikzf2* (H) and *Lbh* (I), are presented in the cells presented in (A). **(J)** Heatmap of the top genes that show significantly different levels between P30_WT IHCs and P30_WT OHCs.

**Supplemental Figure 5.**
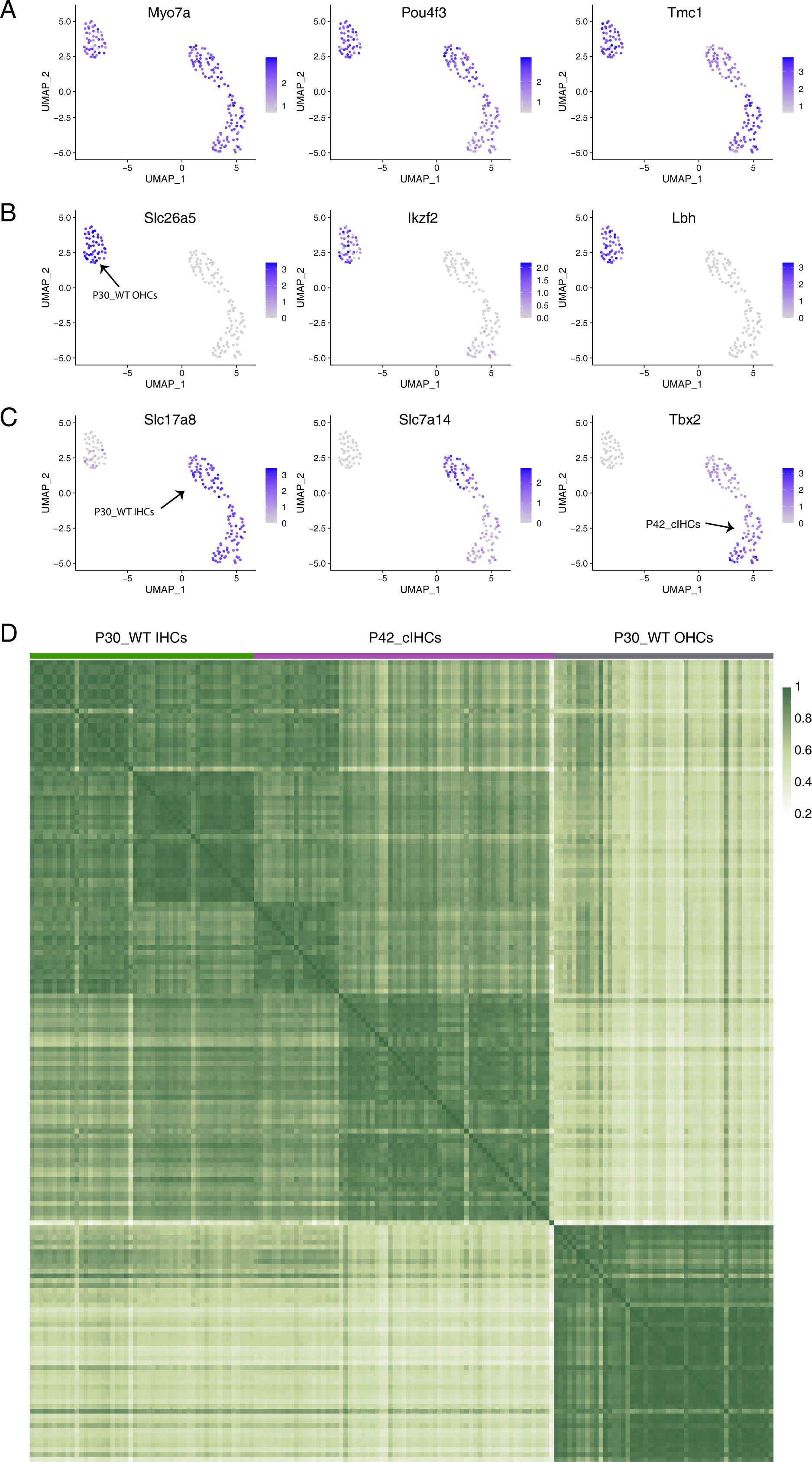
**(A-C)** The cell population (same as in Figure 3A-B), which include P30_WT OHCs, P30_WT IHCs and P42_cIHCs, are plotted with pan HC marker genes *Myo7a*, *Pou4f3* and *Tmc1* (A), OHC specific genes *Slc26a5*, *Ikzf2* and *Lbh* (B), and IHC specific genes *Slc17a8*, *Slc7a14* and *Tbx2* (C). The arrow in (B) points to the P30_WT OHCs. The arrow in (left, C) points to P30_WT IHCs, whereas the arrow in (right, C) points to P42_cIHCs. **(D)** The correlation coefficient analysis of P30_WT OHCs, P30_WT IHCs and P42_cIHCs. The global gene profiles of P42_cIHCs are divergent from the P30_WT OHCs, instead become similar but far from fully equivalent to the P30_WT IHCs.

**Supplemental Figure 6.**
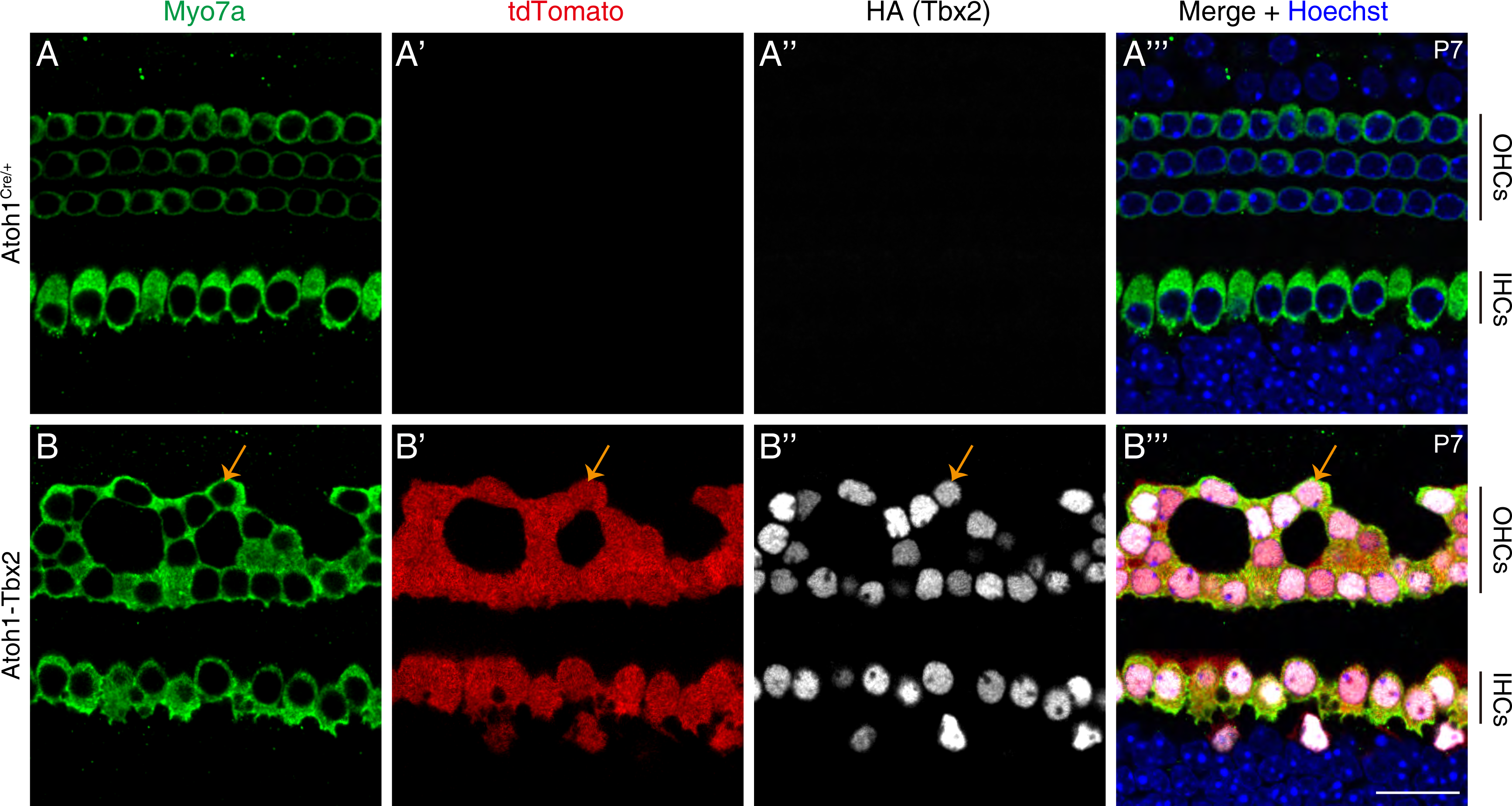
Ectopic Tbx2 is distributed in nuclei of HCs and causes defective cochlear development by P7. Triple staining of Myo7a, tdTomato and HA (Tbx2) in control *Atoh1^Cre/+^*(A-A’’’, n=3) and Atoh1-Tbx2 mice (B-B’’’, n=3) at P7. The tdTomato and HA (Tbx2) expressions are tightly paired and only are detected in Atoh1-Tbx2 mice (B-B’’’). All tdTomato+ cells are HA (Tbx2)+ and vice versa. Orange arrows mark one cIHC that expresses Myo7a, tdTomato and HA (Tbx2). Notably, as expected HA (Tbx2) is distributed in the HC nuclei.

**Supplemental Figure 7.**
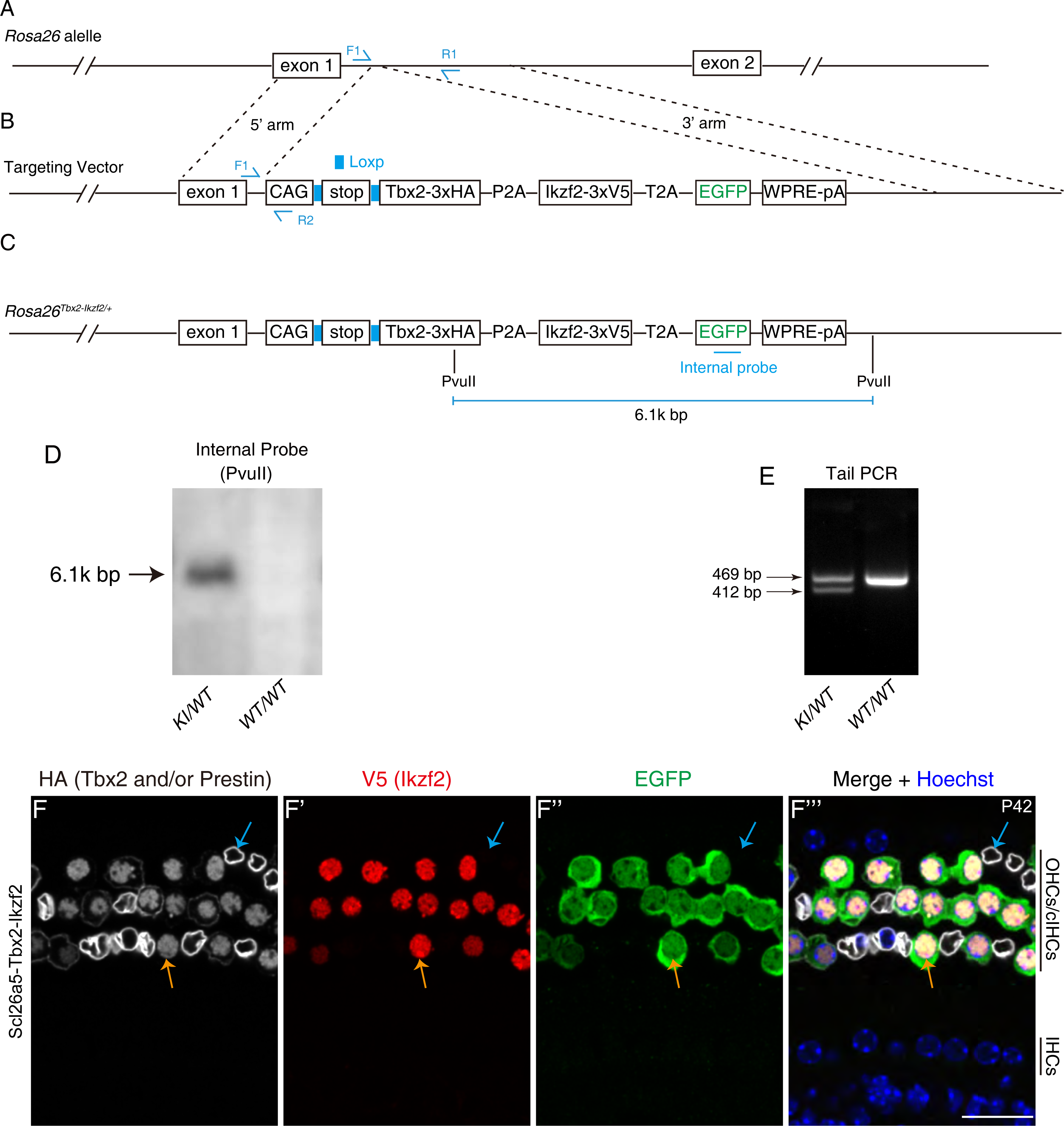
Construction of the *Rosa26*-Loxp-stop-Loxp-Tbx2*3 ◊ HA-P2A-Ikzf2*3 ◊ V5-T2A-EGFP/+ (*Rosa26^Tbx^*^2^*^-Ikzf^*^2^*^/+^*). **(A-C)** In the wild type *Rosa26* locus (A), we insert a DNA fragment between the 5’ and 3’ arms in the targeting vector (B), resulting in the final *Rosa26^Tbx^*^2^*^-Ikzf2/+^* allele (C). **(D)** Southern blot assay with the internal probe. A 6.1k bp band is detected in the heterozygous (KI/WT), but not in the wild type (WT/WT) mice. **(E)** One exampled gel image of tail PCR of heterozygous and wild type mice. **(F-F’’’)** Triple staining of HA (Tbx2) and/or HA (Prestin), V5 (Ikzf2) and EGFP in the Slc26a5-Tbx2-Ikzf2 mice at P42. Prestin is also tagged with HA (membrane) in *Slc26a5^CreER/+^*. Tbx2, Ikzf2 and EGFP expressions are tightly paired. All EGFP+ cells express nucleus HA (Tbx2) and V5 (Ikzf2), and vice versa. The orange arrows mark one EGFP+ cell (or cIHC) that expresses V5 (Ikzf2), and nucleus HA (Tbx2), whereas the blue arrows mark one endogenous OHC that expresses the membrane HA-tagged Prestin but does not express V5 (Ikzf2) and the nucleus HA-tagged Tbx2. Scale bar: 20 μm (F’’’).

